# Altering Mammalian Transcription Networking with Adaadi: An Inhibitor of Atp-Dependent Chromatin Remodeling

**DOI:** 10.1101/723593

**Authors:** Radhakrishnan Rakesh, Saddam Hussain, Kaveri Goel, Soni Sharma, Deepa Bisht, Upasana Bedi Chanana, Joel W. Hockensmith, Rohini Muthuswami

## Abstract

Transcriptional control has been earnestly pursued for the regulation of cellular proliferation associated with cancer progression. The foundational paradigm of targeting transcription factors has yielded exquisite specificity, but many factors cannot yet be targeted. In contrast, targeting epigenetic factors to control chromatin structure and consequential gene expression generally yields more global effects on transcription. Our working paradigm targets neither specific transcription factors nor global epigenetic factors but ATP-dependent chromatin remodeling factors that regulate expression of a limited set of genes. Active DNA-dependent ATPase A Domain inhibitor (ADAADi) synthesized by aminoglycoside phosphotransferases is the first-in-class inhibitor of ATP-dependent chromatin remodeling proteins that targets the ATPase domain of these proteins. Mammalian cells are sensitive to ADAADi but cell lines are variable in their individual responses to the inhibitor. The ADAADi product can be generated from a variety of aminoglycoside substrates with cells showing differential responses to ADAADi depending on the starting aminoglycoside. RNA seq analysis demonstrated that targeting the chromatin remodeling by treatment with a sub-lethal concentration of ADAADi yields alterations to the transcriptional network of the cell. Predominantly, the tumor-promoting genes were repressed while pro-apoptotic and tumor suppressors genes were upregulated on treatment with ADAADi, leading to apoptotic-type cell death. Treatment with ADAADi reversed the EMT process as well as inhibited migration of cells and their colony forming ability. In conjunction with the previous report that treatment with ADAADi regresses tumors in mouse model, this chromatin remodeling inhibitor shows promising anti-tumor properties by targeting the main hallmarks of cancer.

## INTRODUCTION

Chromatin functions as both a structural and regulatory barrier for access to DNA by proteins and enzymes that effect metabolic processes including transcription, recombination, repair and replication, and thus also has an impact on cellular growth and differentiation. Extensive knowledge of the nucleosomal structure of chromatin and the component histones and their post-translational modifiers has yielded a wealth of targets for low molecular weight inhibitors that target chromatin structure. Thus, deregulation of chromatin structure through epigenetic targeting of histone writers, histone erasers and histone readers is a burgeoning field of study with significant implications for the control of cancer cell growth ^1,2^. Available evidence supports that cancer might be treated ^2^ or even prevented ^3^ by chemicals that affect epigenetic players and we have established that cancer progression can be limited through the use of ADAADi, an inhibitor of ATP-dependent chromatin remodeling ^4^.

The body of knowledge regarding ADAADi spans its protein target, the regulation of transcription, the induction of apoptosis and even the clearance of solid tumors from an animal model ^4–7^. Herein we add details regarding the impact of chromatin remodeling on cellular processes with particular interest in aspects of tumorigenesis and its progression.

The six hallmarks that dictate tumorigenesis and its progression are limitless replication, insensitivity to growth-inhibitory signals, self-sufficiency in growth signals, sustained angiogenesis, inhibition of apoptosis, and sustained tissue invasion and metastasis ^8^. It has become increasingly evident that the genes regulating these pathways are regulated by chromatin remodeling, a mechanism involving histone modifications and nucleosome repositioning ^9^.

The ATP-dependent chromatin remodeling proteins have been shown to participate in transcriptional regulation of gene expression by harnessing energy released by ATP hydrolysis to reposition nucleosomes ^10^. These proteins have been classified into 24 sub-families based on phylogenetic analysis ^11^.

Experimental evidence has shown that many of these proteins can function as tumor suppressor genes as well as drivers of cancer ^12^. BRG1 and BRM, transcriptional regulators, have been shown to be mutated or mis-regulated in many cancers ^13–15^. BRG1 has also been shown to cooperate with both BRCA1 and steroid receptors, thus, linking BRG1 to breast cancer ^16,17^. It is also a prognostic indicator for prostate cancer ^4^. Loss of expression of BRG1/BRM has been correlated with poor prognosis in lung cancer ^18^. The NuRD complex contains CHD/Mi-2 as the ATP-dependent chromatin remodeler and though, no mutations have been reported in this protein, yet it is known that the other components of the complex are altered in cancer tissues ^19^. Of the other remodelers, though none of them have been studied for their direct role in tumorigenesis, it is known that many of them participate in DNA repair which is a mechanism intimately linked to cancer development and progression ^20,21^.

Therefore, these proteins have emerged as potential drug targets for cancer chemotherapy. While many inhibitors for the histone modifying enzymes are known, the inhibitors for the ATP-dependent chromatin remodeling proteins are limited in number and scope. Until now only PFI-3 targeting the bromodomain of BRG1 has been reported ^22^. In addition to this inhibitor, we had earlier reported that the enzyme aminoglycoside (neomycin) phosphotransferase catalyzes the formation of a small molecule called Active DNA-dependent ATPase A domain inhibitor (ADAADi) that effectively inhibits the ATPase activity of the ATP-dependent chromatin remodeling proteins ^5^.

ADAADi is generated by the catalytic reaction mediated by aminoglycoside phosphotransferases using aminoglycosides as substrates ^5,6^. Using fluorescence spectroscopy, ADAADi was shown to bind to Active DNA-dependent ATPase A Domain (ADAAD) both in the absence as well as presence of ATP and DNA. This binding to ADAAD occurs via a specific conserved region previously identified as Motif Ia of the SWI2/SNF2 proteins leading to a conformational change in ADAAD that precludes ATP hydrolysis ^6^. Thus, ADAADi has high specificity, targeting the DNA-driven conformational change that effects ATP hydrolysis which is essential for the chromatin remodeling.

The specificity with which ADAADi targets ATP-dependent chromatin remodeling proteins is remarkable since it neither inhibits other types of DNA-dependent ATPases nor DNA-independent ATPases ^5^.

The cellular effects of ADAADi include reduction in the proliferation of mammalian cells lines like PC3, Neuro-2A and MCF-7 implicating a possible role of this molecule as a chemotherapeutic reagent ^4,6,7^. Reinforcement of this concept of regulation of growth signal also emanates from the sensitivity of transcription to ADAADi in the breast cancer cell lines MDA-MB231, MDA-MB-468 and HDQ-P1 with recent studies showing that ADAADi targets BRG1 within the cell and therefore, functions as a potent BRG1 inhibitor ^4,7^.

We have previously established efficacy in the pursuit of BRG1 as a chemotherapeutic target ^4,7^ and in this paper, we endeavor to delve into the complex matrix of effects that stem from the use of different ADAADi species in multiple mammalian cell lines. Each of the latter has its own distinct expression profile and epigenetic state which may account for their differential sensitivity to the ADAADi molecule and the induction of cell death through the apoptotic process. Finally, we also show that ADAADi disrupts the epithelial-to-mesenchymal transition, cell migration and colony formation indicating that the compound has the potential to inhibit some of the hallmark processes of cancer.

## RESULTS

### Cellular responsiveness to ADAADi is complex including time-dependent cellular exposure

ADAADi was discovered as an inhibitor of ADAAD, which is the active ATPase protein domain from the bovine homolog of SMARCAL1 ^5^. Previous studies have shown that ADAADi can be used effectively to kill breast cancer cells ^7^ and prostate cancer cells ^4^. We have expanded these observations to understand the effects of ADAADi on a wide range of cancer cell types and present the findings herein for eight cancer cell lines.

To quantitate the effect of ADAADi on mammalian cells, the time frame required for its action was first investigated. Mammalian cell lines were treated with neomycin-derived ADAADi (ADAADiN) for 24 hr and 48 hr and the viability of the cells was measured. The MTT assay showed that the treatment yielded greater alteration of cellular metabolism at 48 hr as compared to 24 hr (Supplementary Fig. 1). Therefore, all subsequent experiments were done for 48 hr of ADAADi treatment unless otherwise stated. The IC_50,_ in each case, at 48 hr was found to be greater than at 24 hr (Table 1; Supplementary Fig. 1). It should be noted that in case of mechanism-based inhibition, the IC_50_ values depend on the time and therefore, the values are different at 24 and 48 hr ^23^.

**Table 1:** IC_50_ values calculated for different cancer cell lines with ADAADiN (at 24 and 48 hr) and ADAADiK (48 hr). Calculated using: https://www.aatbio.com/tools/ic50-calculator/

| Cells | IC50 (μM) for ADAADiN |  | IC50 (μM) for ADAADiK |
| --- | --- | --- | --- |
|  | 24 hr | 48 hr | 48 hr |
| **A549** | 8.6 | 2.6 | 11.9 |
| **DU145** | 18.4 | 6.9 | 9.8 |
| **PC3** | 20.3 | 10.1 | 8.8 |
| **HepG2** | 9.2 | 4.8 | 12.6 |
| **HeLa** | 24.7 | 10.1 | 10.9 |
| **L132** | 19.5 | 11.3 | 686 |
| **MCF-7** | 18.9 | 3.9 | 379 |
| **MDA-MB-231** | Could not be calculated | 37.9 | 11.9 |
| **HEK293** | Could not be calculated | 24.6 | 6.2 |

### Responsiveness to ADAADi is a function of both the ADAADi derivative and the mammalian cell line

ADAADi can be generated from many different aminoglycosides ^6^. We hypothesized that the ADAADi derived from structurally different starting substrates could have differential effects on mammalian cell lines. To address this postulate, mammalian cancer cell lines were treated with ADAADi generated from neomycin (ADAADiN) or kanamycin (ADAADiK), with the former having four glycosidic rings and the latter having only three glycosidic rings. After 48 hr of treatment, cell lines displayed differential sensitivity to the inhibitors (Supplementary Fig. 1B; Table 1). As an example, the L132 cell line is derived from HeLa cells but this former cell line is extremely resistant to ADAADiK as compared to HeLa even though the cell lines show comparable sensitivity to ADAADiN. Two cell lines (DU145 and PC3) derived from prostate tissue demonstrated comparable sensitivity to ADAADiN but also responded roughly equally to ADAADiK. Unlike the prostate cells, MDA-MB-231 breast cancer cells were most sensitive to ADAADiK while the MCF-7 breast cancer cell line was dramatically more sensitive to ADAADiN. Overall, 7 of the 9 cell lines yielded sensitivity to ADAADiN that was equal to or greater than that of ADAADiK with the exceptions being cell lines HEK293 and MDA-MB-231 which were demonstrably more sensitive to the ADAADiK. However, we were unable to establish any simple rules for the potency of the ADAADi inhibitors either with respect to the origin of the cell line or with respect to type of cell line (Table 1).

### The responsiveness of cells is not dependent on the status of SMARCAL1 or BRG1 alone

To understand the effect of ADAADi at the molecular level, we turned our attention to the expression of SMARCAL1 and BRG1 as we have previously established the sensitivity of SMARCAL1 and BRG1 proteins to ADAADi ^5,6^.

We hypothesized that the responsiveness of the cell to ADAADi would be dependent on the presence or absence of SMARCAL1 and/or BRG1. Thus, a cell line that expresses both SMARCAL1 and BRG1 would be more sensitive as compared to a cell line that expresses either SMARCAL1 or BRG1 while a cell line that expresses neither SMARCAL1 nor BRG1 would be least responsive to ADAADi. To test the hypothesis, the expression of the two proteins in the cancer cell lines was analyzed using western blot (Fig. 1A). The expression of these proteins when compared to responsiveness of cells to ADAADi demonstrated no correlation between the expression of SMARCAL1 or BRG1 and the responsiveness of the cells to ADAADi (Compare Fig. 1A and Table 1). For example, BRG1 is absent in A549 and DU145 cells (Fig 1A as well as shown by Wong et al. ^14^) and therefore, under this hypothesis, these two cells lines would be expected to be resistant to ADAADi as compared to HepG2 and HeLa cell lines, both of which express the protein. However, A549, DU145 and HepG2 show similar response to ADAADi while HeLa is 4-fold more resistant to the inhibitor as compared to A549 (Table 1). Thus, the data suggests that the presence or absence of SMARCAL1 and BRG1 is not the sole determinant of the responsiveness of a cell line to ADAADi

**Figure 1.**
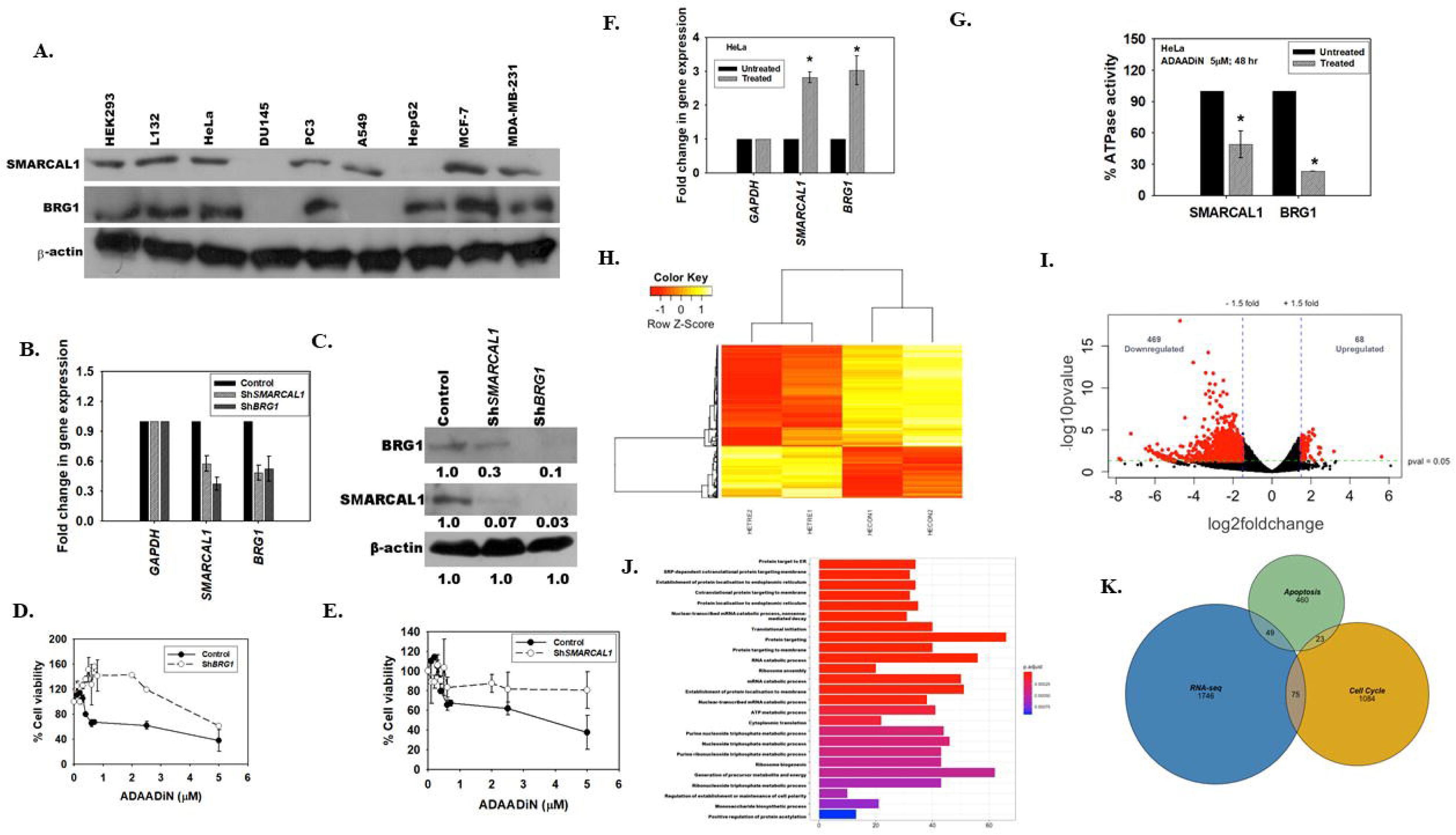
Expression of SMARCAL1 and BRG1 is altered in cancer cell lines. (A). Expression of SMARCAL1 and BRG1 was analyzed by western blot in HEK293, L132, HeLa, DU145, PC3, A549, HepG2, MCF-7, and MDA-MB-231. β-actin was used as loading control. (B). qPCR showing the expression of *BRG1* and *SMARCAL1* in cells after downregulation of these genes in HeLa cells. *GAPDH* was used as internal control. (C). Western blot showing the expression of BRG1 and SMARCAL1 in Sh*BRG1* and Sh*SMARCAL1* HeLa cells. β-actin was used as the internal control and the levels were quantitated using Image J software. (D). Downregulation of *BRG1* renders the HeLa cells resistant to ADAADiN treatment as compared to the control cells. (E). Downregulation of *SMARCAL1* renders the HeLa cells resistant to ADAADiN treatment as compared to the control cells. In these two experiments, *BRG1* or *SMARCAL1* was stably downregulated in HeLa cells and ADAADiN treatment was given for 48 hr. Cell viability was measured using MTT assay. (F). The transcript levels of *SMARCAL1* and *BRG1* in HeLa cells after ADAADiN treatment for 48 hr were measured using qPCR. (G). The ATPase activity of SMARCAL1 and BRG1 was measured in untreated and ADAADiN-treated HeLa cells. In both these experiments, the percent ATPase activity was measure with respect to the untreated control at the respective time points. (H). Heat Map showing the differential expression of genes between untreated and ADAADiN treated HeLa cells. (I). Volcano plot showing the differentially expressed genes between untreated and ADAADiN treated HeLa cells. (J). Gene ontology showing the various classes of genes that are altered on ADAADiN treatment. (K) Intersection between the differentially expressed genes identified by RNA-seq experiment, genes involved in apoptosis and cell cycle. In all these experiments, HeLa cells were treated with 5 μM ADAADiN for 48 hr. The qPCR and ATPase activity data are presented as average ± s.d of three independent experiments. The star indicates p<0.05.

Our second approach was the demonstrable downregulation of BRG1 or SMARCAL1 as a convenient avenue to determine whether the diminished presence of these proteins would alter the response of a cell line towards ADAADi. For this experiment, HeLa cells that express both BRG1 and SMARCAL1 were employed (Fig. 1A). Downregulation was achieved by transfecting HeLa cells with either ShRNA to *BRG1* or to *SMARCAL1* and downregulation was validated by both qPCR and western blot (Fig. 1B and C). Downregulation of *BRG1* or *SMARCAL1* in HeLa cells rendered the cells resistant to ADAADi treatment (Fig. 1D and E) which is congruent with the hypothesis that BRG1 and SMARCAL1 are targets of ADAADi and thus their diminished presence renders the cell resistant to ADAADi. The inference is that these proteins are a primary target of the inhibitor and are necessary for the action of the inhibitor on a cell line. Nevertheless, as shown by the western blotting (Fig. 1A compared with Table 1), the presence/absence of a single ATP-dependent chromatin remodeling factor in the cellular milieu of many related proteins may not be sufficient to predict the response of a cell line to ADAADi.

In all the subsequent experiments we used HeLa cells as they expressed both BRG1 and SMARCAL1. We have treated Hela cells with 5 μM ADAADiN for 48 hr in all the experiments described henceforth.

### The transcriptomic profile of HeLa cells is altered on treatment with ADAADiN

ADAADiN treatment leads to an increase in the transcript levels of SMARCAL1 and BRG1 (Fig. 1F). However, the protein produced is inactive as observed by reduced ATPase activity in the cases of both SMARCAL1 (retains only 46% activity) and BRG1 (retains only 26% activity) in HeLa cells after 48 of treatment (Fig. 1G). The ATPase activity of these proteins is needed for co-regulating transcription ^24–27^. Also, we had previously shown that cell line resistant to ADAADiN showed decreased ATPase activity of SMARCAL1 and BRG1 that manifested in alterations in the transcriptomic profile of the cells^6^. Therefore, we hypothesized that the inactivation of the ATP-dependent chromatin remodeling proteins by ADAADiN would alter the transcriptome of cells. To this end, we treated HeLa cells with ADAADiN for 48 hr and RNA-seq was performed on the isolated transcripts. The RNA-seq data was analyzed using Salmon, DESeq2 taking UCSC known gene file as reference transcriptome to identify the differentially expressed genes. The data was analyzed using two biological replicates, normalized (Supplementary Fig. 2A) and analysis showed good correlation as well as mean-variance relationship between the replicates (Supplementary Fig. 2B and C).

About 800 genes that were involved in protein targeting, metabolic processes and protein translation were found to be differentially regulated (p< 0.05) (Supplementary Excel File 1). The heat map of the top 100 (p<0.05) differentially expressed genes between the untreated and treated samples showed that there was consistency between the replicates and that there are more downregulated genes as compared to upregulated genes on ADAADiN treatment (Fig. 1H). A volcano plot too was generated to observe the overall differences between the untreated and ADAADiN treated samples (Fig. 1I). The plot showed that the expression of 537 genes was altered (p<0.05) by 1.5 fold (Fig. 1I). Gene ontology showed that the differentially expressed genes included regulated protein targeting to ER, ATP metabolic processes, ribosome assembly, and ribosome biogenesis pathways (Fig. 1J). Pathway analysis using KEGG also showed that 49 genes involved in apoptosis and 75 genes involved in cell cycle are differentially expressed on ADAADiN treatment (Fig. 1K). SMARCAL1 and BRG1 were upregulated, as we had found by qPCR (see Fig. 1F), but the change in their expression as well in the expression of other ATP-dependent chromatin remodeling proteins was not statistically significant (Supplementary Excel File 1). Further, the transcripts encoding for *ABCG1*, *ABCB1*, and *ABCG2* proteins (multi-drug resistance pumps) were found to be unaltered in the case of HeLa cells. This is consistent with the lack of change in basal transcription previously reported for MDA-MB-231 cells but which is unlike the effect of ADAADiN upon transcription of these genes when induced by a variety of chemotherapeutic reagents wherein ADAADiN treatment prevents the upregulation of these pumps ^7^. Thus, we infer that basal regulation of ABC cassette genes by ATP-dependent chromatin remodelers may employ different mechanisms than the upregulation of the same genes.

As RNA-seq data showed some of the splicing factors are altered on ADAADiN treatment, we inquired whether ADAADiN treatment is causing the change in transcript usage after the treatment. For this purpose, the data was also analyzed using the GENCODE reference transcriptome and nearly 60 percent correlation of differentially expressed genes (log2foldchange>1.3) was observed between the two analyses. The differential transcript usage on RNA-seq data obtained when GENECODE reference transcriptome was used was analyzed by DTUrnaseq-DRIMSeq package. We used GENECODE data for the splice variant usage analysis because it contains considerably more annotated and predicted transcripts annotation than the known gene reference transcriptome. Our analysis showed that about 900 genes were switching to alternate transcripts after ADAADiN treatment (Supplementary Excel File 2).

To identify the genome wide occupancy of SMARCAL1 and BRG1 with the implication of a better understanding of the transcriptome regulation by these ATP-dependent chromatin remodeling proteins, ChIP-seq was performed using antibodies against SMARCAL1 and BRG1.

We employed, untreated HeLa cells. Unsupervised k-mean clustering (k=5) of ChIP-seq peaks showed that the two proteins are present on the transcription start sites (TSS) (Fig. 2A) and their peak overlap showed that these proteins co-occupy 8755 sites on the genome (Fig. 2B; Supplementary Excel File 3 and 4). Previously, we had shown that SMARCAL1 was present on *BRG1* promoter and BRG1 was present on *SMARCAL1* promoter ^24^. The ChIP-seq data validated our experimental results (Fig. 2C; Supplementary Excel File 3 and 4) as SMARCAL1 and BRG1 were found to be present on *SMARCAL1* and *BRG1* genes. SMARCAL1 and BRG1 were also found on the promoter of *ATM* gene (Fig. 2C), once again validating the experimental results obtained in previous studies ^26^. Finally, we analyzed the presence of these two proteins on ABC transporters as previously BRG1 has been shown to regulate the expression of these genes in breast cancer cells ^7^. The ChIP seq data showed that both BRG1 and SMARCAL1 are also present on *ABCG1*, *ABCB1*, and *ABCG2* genes (Supplementary Fig. 2D), even though their expression does not alter on ADAADiN treatment (Supplementary Excel File 1). Interestingly, peak analysis on the genome browser showed that SMARCAL1 and BRG1 were also found to occupy intronic regions specially the first intron of several genes.

**Figure 2.**
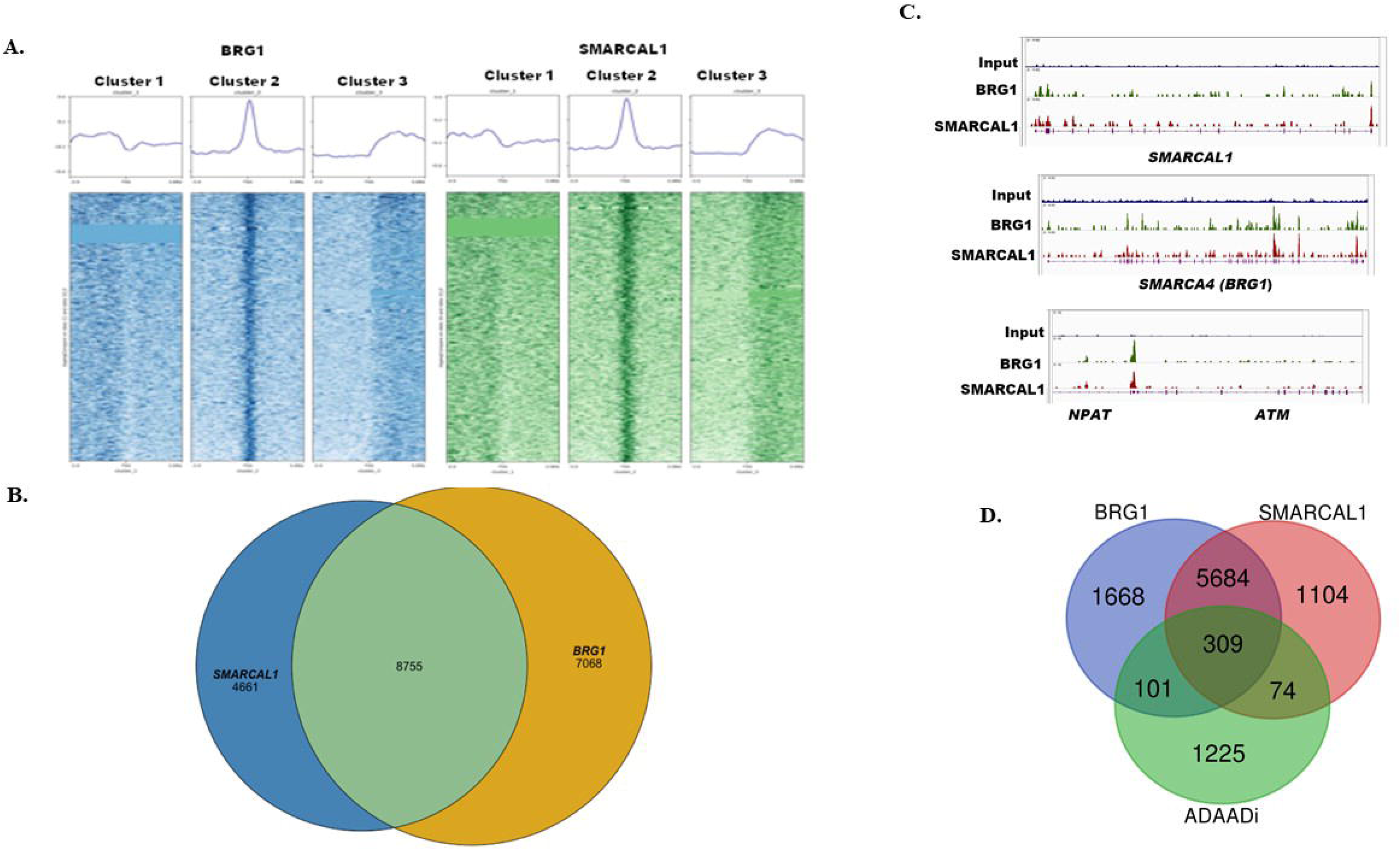
The genome-wide occupancy of SMARCAL1 and BRG1 was studied using ChIP-seq. (A) Unsupervised k-mean (k=5) clustering of chip peaks (BRG1-blue, SMARCAL1-green). SMARCAL1 and BRG1 shows highest peak occupancy over TSS on cluster 2. There was no binding over TSS on cluster 3 and 4. (B) ChIP peak intersection between SMARCAL1 and BRG1. (C) Representative ChIP-seq peaks visualized using IGV genome browser showing occupancy of SMARCAL1 and BRG1on *SMARCAL1*, *BRG1*, and *ATM* genes. (D) Intersection among genes occupied by SMARCAL1, BRG1 and differentially expressed genes (p<0.1) after ADAADiN treatment.

Using a less stringent condition p<0.1 led us to identify 1710 differentially regulated genes of which 484 genes (28.3% genes) were found to be regulated by SMARCAL1 and BRG1 either alone or by a combination of the two proteins (Fig. 2D). The genes co-regulated by SMARCAL1 and BRG1 include *ROCK2* and *EGF* (Fig. 3A; Supplementary Excel File 3 and 4). ROCK2 is a Rho-associate protein kinase 2 involved in many cellular pathways including apoptosis ^28^, while EGF is a growth factor.

**Figure 3.**
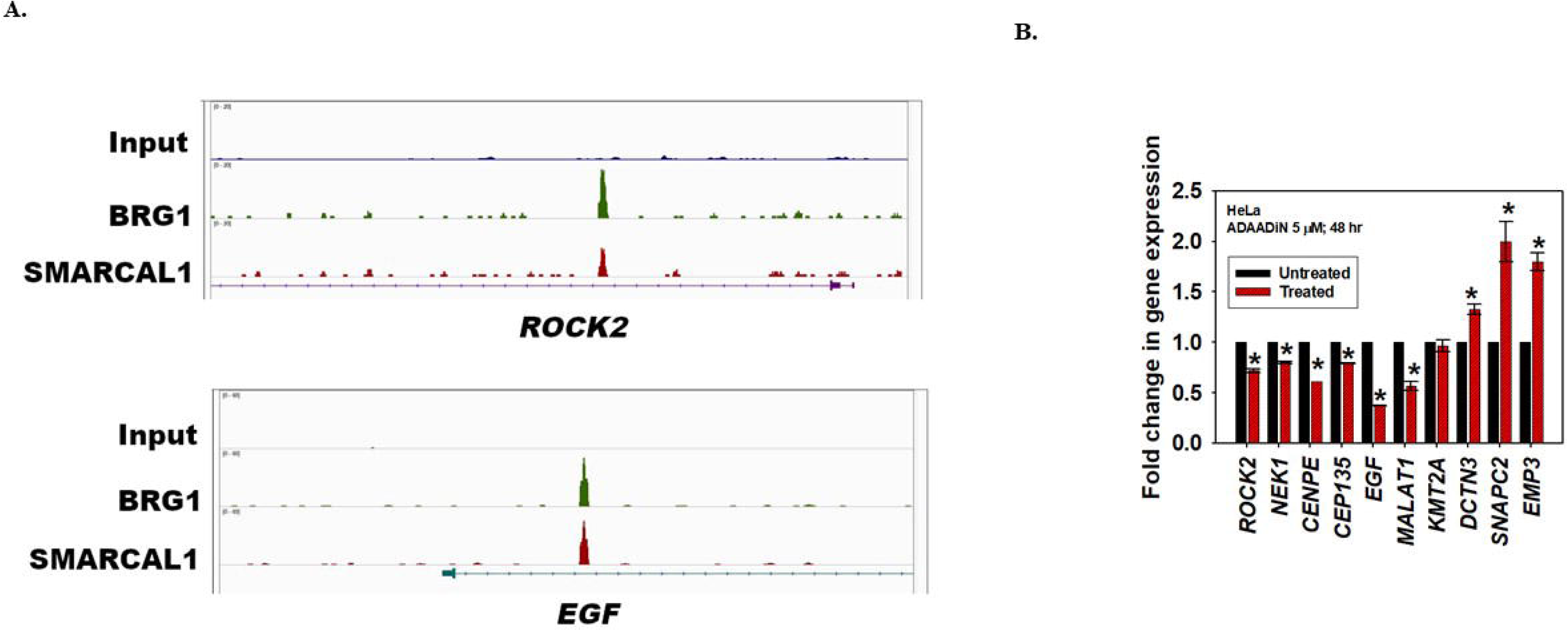
ADAADiN treatment alters gene expression in HeLa cells. (A) IGV genome browser track showing SMARCAL1 and BRG1 binding on *ROCK2*, and *EGF* genes. (B). qPCR validation of RNA-seq data for selected genes. The data is presented as average ± s.e.m of three independent experiments. The star indicates p<0.05.

To validate the ChIP seq data obtained in our experiments, we compared it with the dataset for BRG1 available in public domain and confirmed that BRG1 in both experiments was indeed present on *SMARCAL1*, *BRG1*, *ATM*, *ROCK2*, and *EGF* (Supplementary Fig. 3)

Finally, we sought to experimentally validate the RNA-seq data using qPCR. We picked genes where the occupancy of SMARCAL1 and BRG1 was predicted by ChIP-seq data (*ROCK2, EGF, CENPE*, and *KMT2A*) as well as those genes where the occupancy of these two proteins was not shown by ChIP-seq data (*DCTN3, SNAPC2*, *MALAT1*, *CEP135* and *EMP3*). Experimental results showed that the expression of *ROCK2*, *CENPE*, *CEP135, MALAT1, EGF* was downregulated while the expression of *DCTN3*, *SNAPC2*, and *EMP3*, was upregulated correlating with the RNA-seq data (Fig. 3B). The expression of *KMT2A* was unaltered though the RNA-seq data indicated that the expression of this gene was significantly upregulated (Fig. 3B). Thus, of the 9 genes, we were able to validate the RNA-seq data for 8 genes by experimentation.

### ADAADiN treatment induces DNA damage in a cell type specific manner

RNA-seq data as well as qPCR confirmed that the expression of *ATM*, the transducer of DNA damage signal, was downregulated (p<0.05) (Fig. 4A) in HeLa cells. Since ATM as well as many ATP-dependent chromatin remodeling proteins directly mediate DNA damage response/repair ^29^, we hypothesized that downregulation of ATM expression coupled with inactivation of ATP-dependent chromatin remodeling proteins should result increased DNA damage. To test the hypothesis, we investigated the formation of γH2AX ^30,31^ and 53BP1 foci ^32^ and found to our surprise, that in HeLa cells, neither γH2AX nor 53BP1 foci were observed indicating that DNA damage is not induced in these cells (Fig. 4B). This was further confirmed by comet assay (Fig. 4C and D). To confirm that DNA damage is indeed not induced, we studied the effect of ADAADiN in DU145 cells that do not express SMARCAL1 and BRG1. In these cells, *ATM*, was found to be upregulated on ADAADiN treatment (Fig. 4A). Further in these cells, γH2AX but not 53BP1 foci was observed on ADAADiN treatment (Fig. 4E). Comet assay confirmed that ADAADiN treatment does indeed result in the DNA damage (Fig. 4F and G), suggesting that the ability of the inhibitor to induce DNA damage varies with cell line and is not a factor of expression of DNA damage response genes.

**Figure 4.**
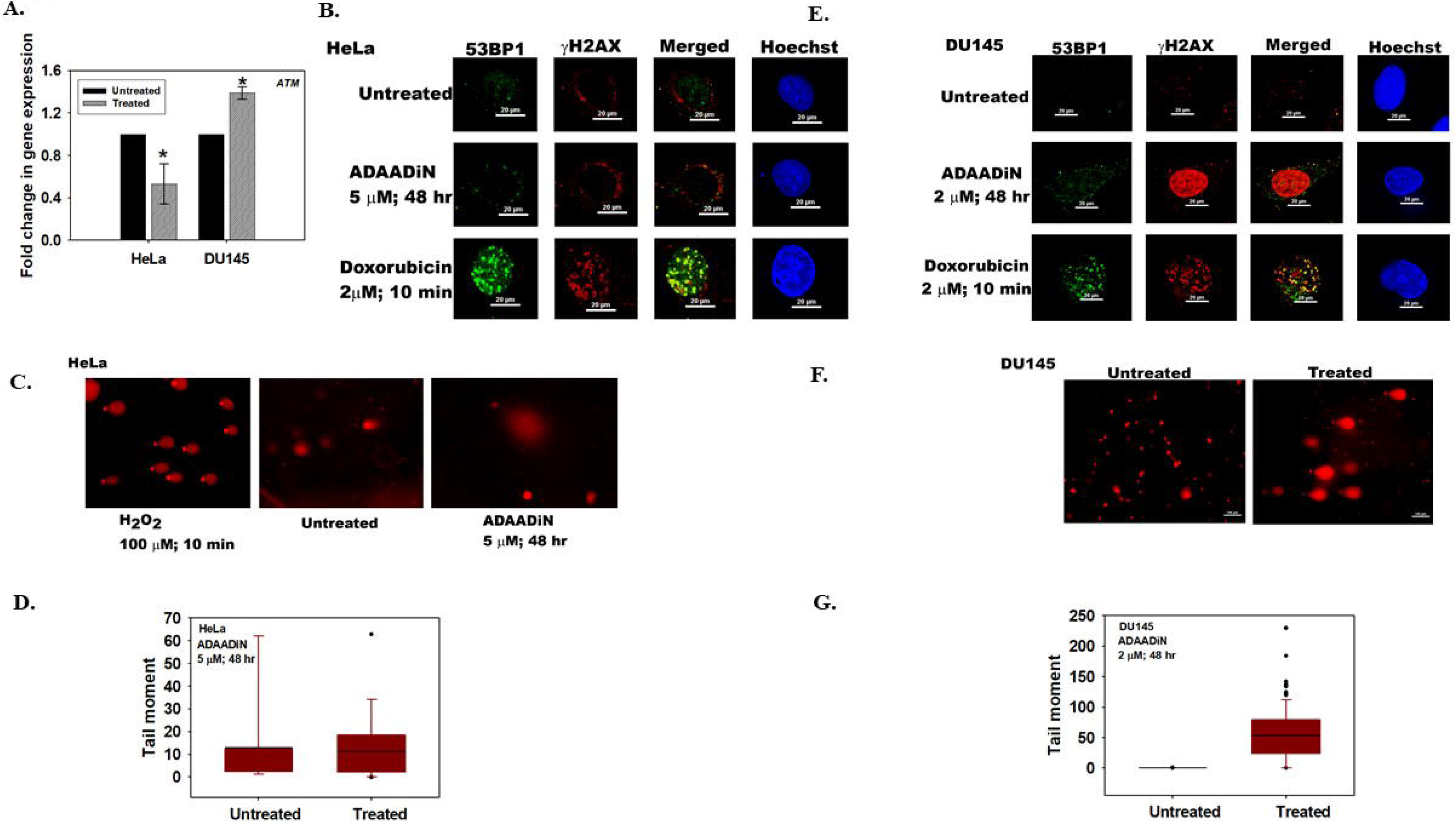
ADAADiN treatment alters the expression of DNA-damage response genes. (A). The transcript level of *ATM* was estimated using qPCR in HeLa and DU145 cells after treatment with ADAADiN. The data is presented is average ± sd of three independent experiments. The star indicates p<0.05. (B). Formation of γH2AX and 53BP1 foci was assessed in HeLa cells after treatment with 5 μM ADAADiN for 48 hr using immunofluorescence. For this experiment, 2 μM doxorubicin treatment for 10 min was used as a positive control. (C). Comet assay in the presence of 100 μM H_2_O_2_ treatment for 10 minutes (positive control), untreated, and ADAADiN treated (48 hr) HeLa cells. (D). Quantitation of tail moment in untreated and ADAADiN treated HeLa samples. (E). Formation of γH2AX and 53BP1 foci was assessed in DU145 cells after treatment with ADAADiN for 48 hr using immunofluorescence. (E). Comet assay performed in untreated and ADAADiN treated (48 hr) DU145 cells. (F). Quantitation of tail moment in untreated and ADAADiN treated DU145 samples.

### ADAADiN treatment results in apoptosis

Previously, we had shown that apoptosis is induced in PC3 cells on ADAADiN treatment ^4^. The HeLa RNA-seq data revealed that 49 genes involved in apoptosis are differentially expressed on ADAADiN treatment (Fig. 1K). Of these *EMP3*^33^ was significantly upregulated while*, ATM* ^34^, and *BRCA1*^35^ were significantly downregulated (p<0.05) and the downregulation was validated by qPCR showed in ADAADiN-treated HeLa samples (Fig. 3B, 4A and 5A respectively). We confirmed apoptotic-type of cell death in both HeLa and DU145 as the two cell types showed differences in DNA damage induction. ADAADiN treatment resulted in an increase in AnnexinV positive cells both in HeLa and DU145 cells as compared to the untreated control (Fig 5B-E). Apoptosis in ADAADiN treated cells was also quantitated using acridine orange/ethidium bromide staining as this method allows for discrimination between live, apoptotic (early and late) and necrotic cells ^36,37^. The results indicated that the number of apoptotic cells increased after ADAADiN treatment as compared to the untreated control in both HeLa and DU145 cells (Fig. 5F-H). Concomitantly, there was a decrease in the number of viable cells after ADAADiN treatment (Fig. 5G and H). Taken together, these results indicate that the cancer cells undergo apoptosis on ADAADiN treatment irrespective of the cell line and whether DNA damage is induced^38^ or not ^39^.

**Figure 5.**
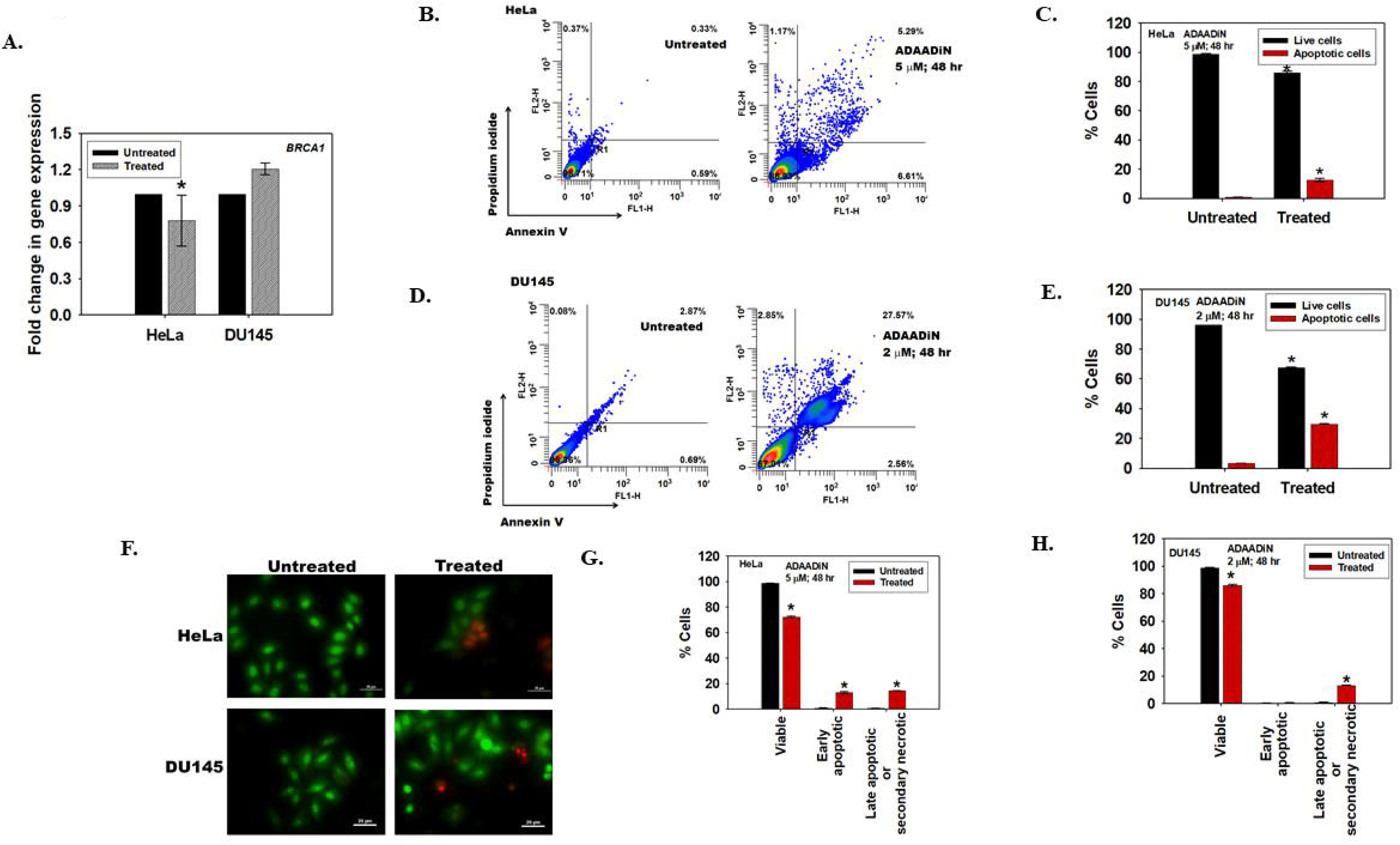
ADAADiN treatment induces apoptosis. (A). The transcript level of *BRCA1* was measured using qPCR in HeLa cells after treatment with 5 μM ADAADiN and in DU145 after treatment with 2 μM ADAADiN for 48 hr. (B). FITC-Annexin V and PI staining in HeLa cells was monitored using FACS in untreated and 5 μM ADAADiN treated HeLa cells. (C). Quantitation of the live and apoptotic cells from FACS analysis for HeLa cells. (D). FITC-Annexin V and PI staining in DU145 cells was monitored using FACS in untreated and 2 μM ADAADiN treated cells. (E). Quantitation of the live and apoptotic cells from FACS analysis for DU145 cells. The data in case of both cell lines is presented as average ± s.d. of three independent experiments. (F). Confocal images obtained after staining untreated and ADAADiN treated HeLa and DU145 cells with acridine orange and ethidium bromide. Green stained cells are viable cells while orange/red cells indicate non-viable or apoptotic cells. (G). The number of viable and apoptotic cells were counted in the untreated and treated HeLa cells. (H). The number of viable and apoptotic cells were counted in the untreated and treated DU145 cells. The results are presented as average ± s.d. of two independent experiments. In each experiment, more than 100 cells were counted. The qPCR data is presented as average ± s.d of three independent experiments. Statistical analysis for qPCR experiments was done using Sigma-plot and the star indicates significance at p<0.05.

### ADAADiN treatment pushes the cell towards epithelial phenotype

With apoptosis established as the mechanism of cell death caused by ADAADiN, we turned our attention to another hallmark of cancer-migration and invasion- and asked whether ADAADiN can block the invasive property of cancer cells ^8^.

The process of epithelial-to-mesenchymal transition (EMT) is known to make the cells shed their epithelial characteristics to acquire more mesenchymal phenotype which is metastatic and renders resistance to chemotherapeutic drugs. Upon examining the RNA-seq data it was found that the genes (*VIM, FN1, CDH1*, and *CDH2*) associated with EMT were not significantly differentially expressed on ADAADiN treatment. However, ChIP-seq data showed that SMARCAL1 and BRG1 are present on *VIM*, *CDH1*, and *CDH2* (Supplementary Excel File 3-5). It has also been shown previously that ADAADiN treatment makes the metastatic cells, MDA-MB-231 more sensitive to chemotherapy ^7^. One plausible explanation for this could be that SMARCAL1 and BRG1 do not regulate the basal transcription of EMT genes and act only upon EMT inducing signal. Therefore, to investigate the role of ADAADiN in the process of EMT, HeLa and DU145 cells were treated with EMT inducing agent TGF-β1 alone or with ADAADiN for 48 hr. Treatment with TGF-β1 alone decreased the transcript levels of the pro-epithelial marker, E-cadherin (CDH1), in both HeLa and DU145 cells (Fig. 6A and B); however, the addition of ADAADiN along with TGF-β1 resulted in the increased expression of *CDH1* (Fig. 6A and B). Fibronectin (FN1), N-cadherin (CDH2), Vimentin (Vim) are pro-mesenchymal whose increased expression has been correlated with EMT ^40^. In DU145 cells, the expression of *FN1* and *CDH2* decreased while in HeLa the expression of *VIM* decreased when these cell lines were treated with ADAADiN along with TGF-β1 as compared to TGF-β1 alone (Fig. 6A and B). EMT is also characterized with decreased expression of Claudin-1 ^40^; western blots confirmed that the expression of Claudin-1 as well N-cadherin returned back to normal levels when the cells were treated with ADAADi and TGF-β1 (Fig. 6C and D), suggesting that ADAADiN treatment does alter the expression of proteins when EMT is induced with TGF-β1; however, the inhibitor has no effect on the basal levels expression of the genes associated with the process.

**Figure 6.**
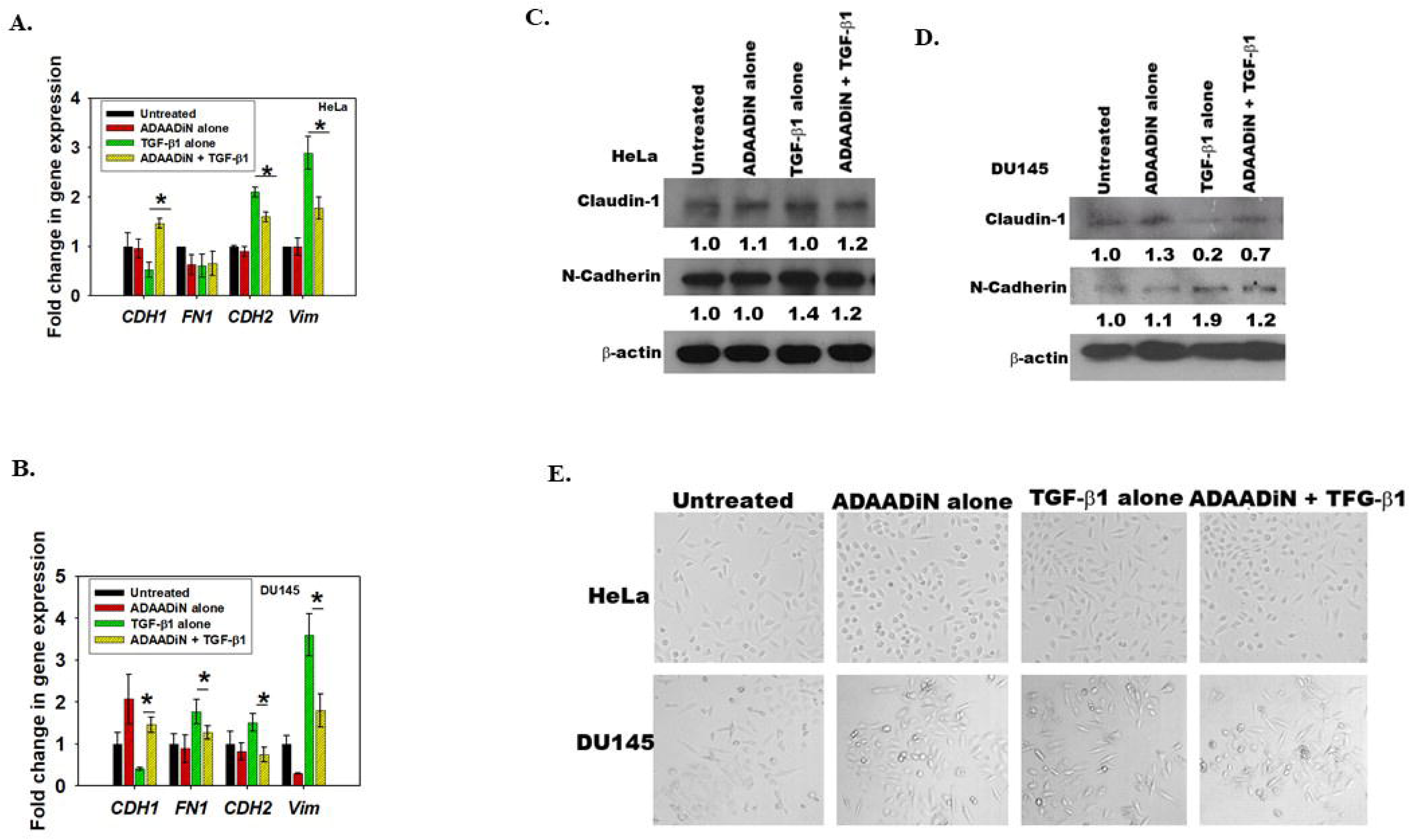
ADAADi treatment pushes the cell towards epithelial phenotype. qPCR analysis of *CDH1*, *FN1*, *CDH2*, and *Vim* expression in (A). HeLa cells; (B) DU145 cells. Western blot analysis for the expression of Claudin-1 and N-cadherin in (C). HeLa cells; (D) DU145 cells. (E). The morphology of HeLa and DU145 cells was analyzed using Nikon TiS microscope in untreated, ADAADiN alone, TGF-β1 alone, and ADAADiN along with TGF-β1 treated cells. In these experiments, cells were treated with ADAADiN or TGF-β1 or ADAADiN and TGF-β1 for 48 hr prior to analysis. *GAPDH* was used as internal control for qPCR experiments and β-actin was used as internal control of western blot analysis. The western blots were quantitated using Image J software. The qPCR data is presented as average ± s.d of two independent experiments. Star indicates statistical significance at p<0.05.

The morphology of the cells was also studied under microscope and the epithelial phenotype of the cells in the presence of ADAADiN and TGF-β1 as was compared to TGF-β alone was evident when analyzed using microscope (Fig. 6E).

Taken together, these results led us to conclude that ADAADiN treatment of cancer cells pushes them towards epithelial phenotype, thus, preventing or delaying TGF-β1-induced transition into mesenchymal phenotype.

### ADAADiN treatment inhibits migration and invasion of cancer cells

Since ADAADiN was found to inhibit the process of EMT which makes the cells more invasive and migratory, we next investigated the levels of matrix metalloproteinases as their role in cancer metastasis has been well-established ^41^. These proteins are secreted into the extracellular milieu where they promote cancer cell migration by degrading the physical barriers. Both MMP-2 and MMP-9 expression has been shown to be associated with tumor aggressiveness and poor prognosis ^41,42^. ChIP-seq data showed that BRG1 and SMARCAL1 are present on *MMP-2* and *MMP-9* promoter (Supplementary Excel File 3-4). ADAADiN treatment resulted in decreased expression of *MMP-2* and *MMP-9* transcripts in both HeLa and DU145 cells (Fig. 7A). RNA-seq data too confirmed that these transcripts are downregulated but not in a statistically significant manner (Supplementary Excel File 1). Further, pro-MMP-2 secretion in HeLa cells and pro-MMP-9 secretion in DU145 cells decreased on ADAADi treatment as observed by zymography assay (Fig. 7B-E). Concomitantly, wound healing assay in both HeLa and DU145 cells showed that the ability to heal the wound was considerably retarded on ADAADiN treatment (Fig. 7F-I).

**Figure 7.**
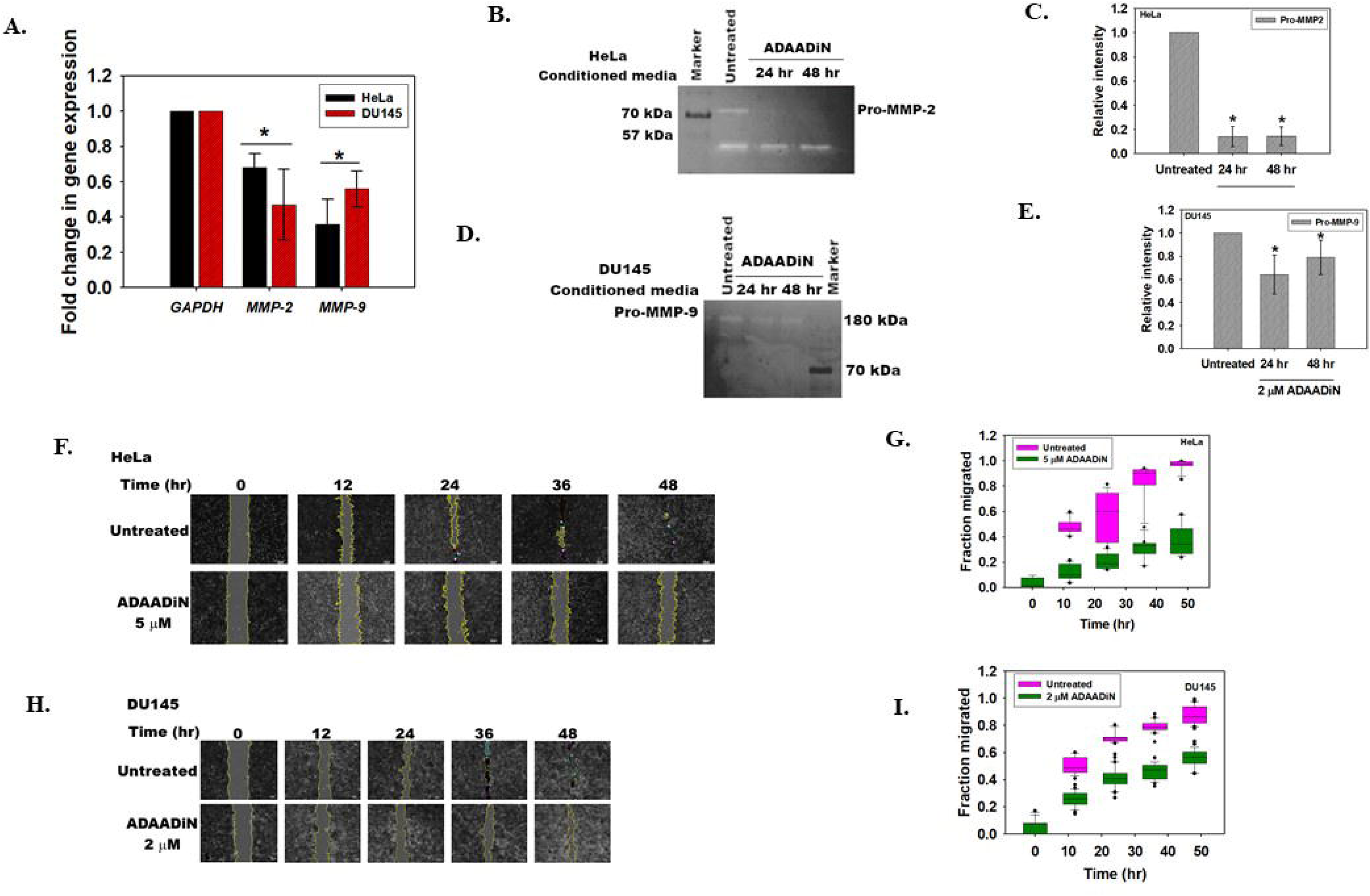
ADAADiN treatment inhibits migration and invasion of cells. (A). qPCR analysis of *MMP-2* and *MMP-9* expression in untreated and ADAADiN treated HeLa and DU145 cells. (B). Zymography assay showing the secretion of Pro-MMP-2 in the media by HeLa cells. (C). Quantitation of Pro-MMP2 was done using Image J software. The data was normalized with respect to untreated control and is presented as average ± s.d. of two independent experiments. (D). Zymography assay showing the secretion of Pro-MMP-9 in the media by DU145 cells. (E). Quantitation of Pro-MMP9 was done using Image J software. The data was normalized with respect to untreated control and is presented as average ± standard deviations of two independent experiments. (F). Image analysis of the wound assay captured at different time points after induction of gap in monolayer of HeLa cells. (G). Quantitation of the migration of HeLa cells in the absence and presence of ADAADiN as a function of time. The data was normalized with respect to untreated control and is presented as average ± s.d. of three independent experiments. (H). Image analysis of the wound assay captured at different time points after induction of gap in monolayer of DU145 cells. (I). Quantitation of the migration of DU145 cells in the absence and presence of ADAADiN as a function of time. The data was normalized with respect to untreated control and is presented as average ± s.d. of three independent experiments. The qPCR data is presented as average ± s.d. of three independent experiments and the star indicates significance at p<0.05.

Taken together, it leads us to conclude that ADAADiN treatment has the potential to retard the invasiveness of cancer cells by altering the expression of metalloproteases and EMT genes.

### ADAADiN inhibits the clonogenicity of the cancer cells

Limitless replication is yet another hallmark of cancer cells. We hypothesized that ADAADiN should be able to inhibit the ability of cancer cells to replicate indefinitely. To test this hypothesis, colony forming assay was performed as explained in the Methods section. Two conditions were used to analyze whether ADAADiN can inhibit colony formation. In the first condition, ADAADiN was added after seeding the cells in the agar containing media. The colony formation ability was determined in the presence of ADAADiN for 12 days. In the second condition, cells were pre-treated with ADAADiN for 24 hr. These cells were seeded in the agar containing media and were allowed to grow in DMEM media that lacked the inhibitor (termed as “pre-treated and discontinued”).

ADAADiN treatment reduced the colony forming ability of both HeLa and DU145 cells as compared to the untreated control as measured by relative area of the colony and Hoechst staining (Fig. 8A-D). Interestingly, pre-treated and discontinued condition also resulted in decreased colony forming ability, suggesting that pre-treatment of cells with ADAADiN is sufficient to block the ability of the cancer cells to replicate indefinitely (Fig. 8A-D).

**Figure 8.**
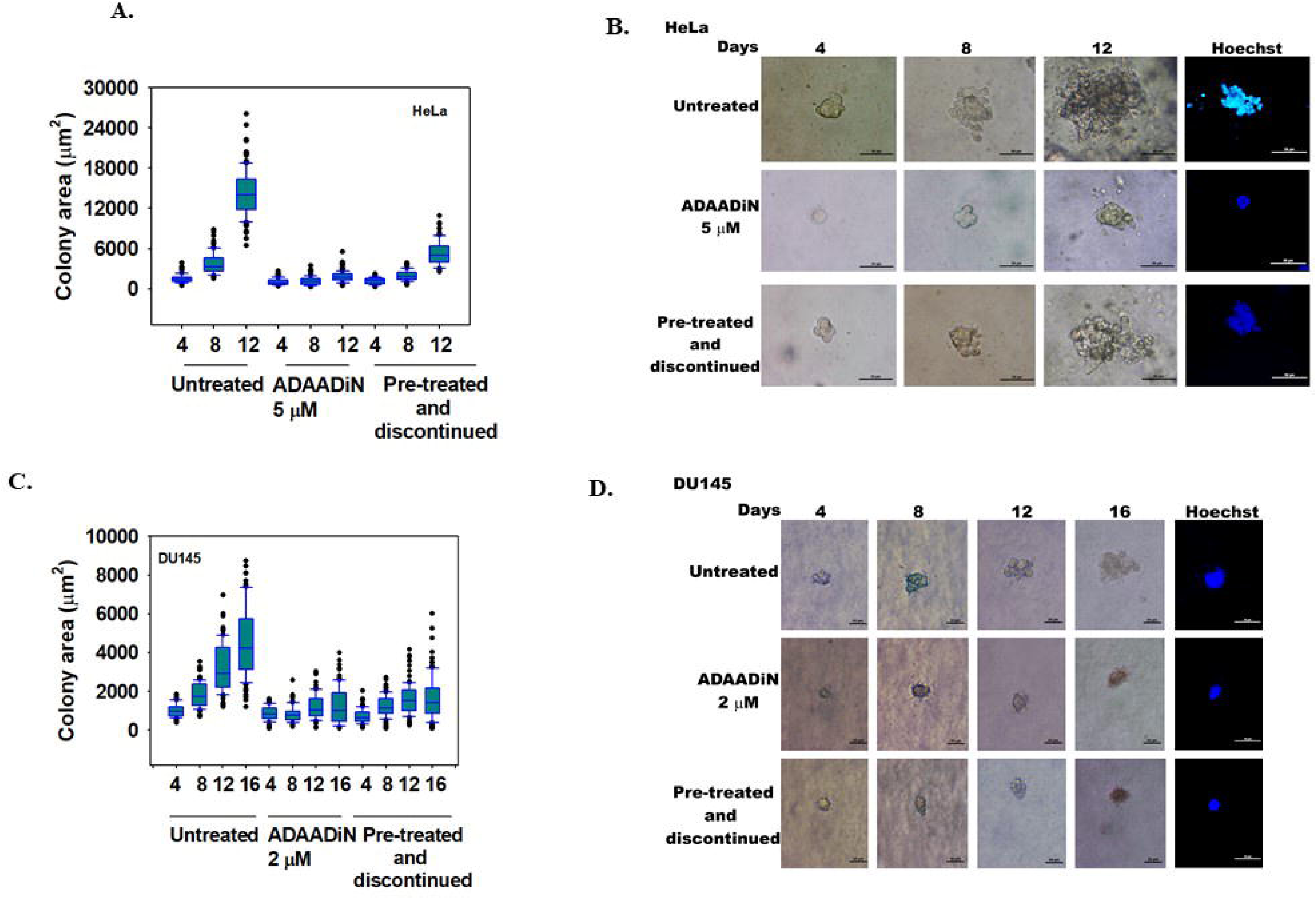
ADAADiN treatment inhibits the colony formation ability. (A). Area (μm^2^) of the HeLa colonies calculated as a function of time in the absence and presence of ADAADiN. (B). Colony formation monitored in untreated, ADAADiN treated, and ADAADiN pre-treated followed by discontinuation of the inhibitor in HeLa cells. On the 12^th^ day, the colonies were stained with Hoechst. Images were taken using Nikon TiS microscope. (C). Area (μm^2^) of the DU145 colonies calculated as a function of time in the absence and presence of ADAADiN. (D). Colony formation monitored in untreated, ADAADiN treated, and ADAADiN pre-treated followed by discontinuation of the inhibitor in DU145 cells. On the 16^th^ day, the colonies were stained with Hoechst. Images were taken using Nikon TiS microscope.

## DISCUSSION

The ATP-dependent chromatin remodeling proteins function as co-regulators of transcription in eukaryotic cells. By coupling ATP hydrolysis to nucleosome repositioning, eviction or histone variant exchange, these proteins are able to regulate the expression of a subset of genes, and therefore, the cellular function ^10^. Thus, it follows that an inhibitor specific to ATP-dependent chromatin remodeling might impact a limited portion of the transcriptome thereby changing the expression of factors that impinge upon the cell fate.

ADAADi is a specific inhibitor of ATP-dependent chromatin remodeling and our results reveal its impact on mammalian cell fate, ranging from survivable alterations in cellular transcription at intermediate levels of ADAADi to induction of apoptosis and cell death at higher levels. In particular, our results reveal complex responses likely related to the hundreds of genes regulated by the target proteins leading to the suggestion that the response a cell line mounts upon ADAADi exposure is probably a complex network of expression of the target proteins, the structure of the ADAADi molecule and factors affecting the IC50 of the inhibitor such as cellular uptake mechanisms and binding affinity of the inhibitor to the target protein. Deciphering the contribution of each of these factors will be required to further understand the responses exhibited by different cancer cell lines towards ADAADi.

By inhibiting the ATPase activity of ATP-dependent chromatin remodeling proteins and therefore the subsequent dependent functions of the enzyme or its complex(es), ADAADi changes the transcriptome and thus, alters the fate of the cancer cell. In general, a change in the transcriptome of the cell in response to chemotherapeutic agents is not surprising as it has been reported for many agents including doxorubicin ^43,44^ and cisplatin ^45^. However, the use of ADAADi contributes new perspectives for consideration. For instance, we have presented multiple examples of genes for which ADAADi has no effect on the basal transcription but whose regulation by various secondary methods are mitigated by the presence of the ADAADi. Thus, the control of ATP-dependent remodeling provides a plausible avenue for control of processes such as cell migration and EMT, as well as transcriptome alterations impacting a wide range of properties of cancer cells. The alteration in transcriptome on ADAADi treatment induces apoptosis, inhibits metastasis, and inhibits replication ability of cancer cells. Our work lays the preliminary foundation demonstrating that ADAADi treatment of cells provides a mechanism for regulation of a discrete subset of all genes, as well as regulation of induced alterations in the transcriptional network of mammalian cells.

The RNA-seq data also revealed that the subset of ADAADi-impacted genes included both protein-coding as well as non-coding transcripts. Moreover, within the subset of protein-coding genes there are a few genes that are associated with cancer progression that show distinctive downregulation by ADAADi treatment, including *EGF*, *CENPE* ^46^, and *ROCK2*^28^. Similarly, non-coding RNAs contributing to the cancer progression like *MALAT1*^47^ is downregulated on ADAADi treatment. When these observations are coupled with the alterations in the expression of DNA-damage response proteins such as *ATM* by ADAADi, the potential for the use of ADAADi in the treatment of cancer becomes more apparent. Nevertheless, there is no theoretical reason why the targets of ADAADi should behave identically in every cell line and even though the expression of DNA damage response genes was altered in HeLa cell line DNA damage *per se* could not be detected either by γH2AX staining or by comet assay. In contrast, in DU145 cells, DNA damage could be detected both by γH2AX staining and comet assay. Such differences in cellular responses may ultimately play a role in the selection of the ADAADi species to be used in studies or even treatment of specific cancer cell lines.

Irrespective of whether DNA damage is induced or not, treatment with ADAADi induces apoptosis as confirmed by Annexin V staining and RNA-seq data that shows 49 genes involved in apoptosis are differentially regulated. Notably, both *ATM* and *BRCA1* are downregulated leading to induction of apoptosis while we were surprised to find that ADAADi treatment does not alter the expression of caspases significantly.

Finally, a major property of cancer cells is their ability to migrate and thus, metastasize to new sites. Whether the effect of ADAADi is interpreted from the perspective of rewriting portions of the transcriptome or the induction of the expression of pro-epithelial genes, ADAADi treatment demonstrably blocks EMT, migration, and invasiveness of cancer cells.

Thus, through its targeting of ATP-dependent remodeling, ADAADi is unlike any other known epigenetic inhibitor. While less specific than targeting a specific transcription factor such as a steroid receptor, ADAADi nevertheless demonstrates sufficient specificity to avoid broad global effects that would lead to general cell death. The demonstrable effects that we document herein illustrate a great chemotherapeutic potential of this novel class of drugs that warrants additional development and prompt evaluation both as an individual and a combinatorial chemotherapeutic approach to cancer treatment.

## METHODS

### Chemicals

All chemicals were of analytical grade and were purchased from Sigma-Aldrich (USA), Qualigens (India), Merck (India), Himedia (India) and SRL (India). LB media was purchased from Himedia (India). Ni^+2^-NTA resin, Protein G beads and Taq polymerase were purchased from Merck-Millipore (USA). cDNA synthesis kit was purchased from Thermo-Fisher (USA). Biorex-70 resin was purchased from Bio-Rad (USA). Cell culture chemicals were purchased from Sigma-Aldrich (USA) while cell culture plasticware was from Corning (USA). SYBR Green mix was purchased from Kapa Biosystems (USA).

### Antibodies

SMARCAL1 antibody was raised against N-terminus HARP domain by Merck (India) (Catalog # 106014). BRG1 (Catalog# B8184) and β-actin (Catalog# A1978) antibodies were purchased from Sigma-Aldrich (USA).

### Primers

The qPCR primers were synthesized by Sigma-Aldrich (USA) and the list of primers used in this study is provided in Supplementary Tables 1 and 2.

### Synthesis and purification of ADAADi

ADAADi was synthesized and purified as explained previously in Dutta *et al.*^6^

### Cell lines

All cell lines used in this study were purchased from NCCS, Pune, India and maintained in Dulbecco*’*s Modified Eagle Medium (DMEM) supplemented with 10% fetal bovine serum (FBS), 1% penicillin-streptomycin-amphotericin cocktail at 37°C in the presence of 5% CO_2_.

### Cell viability assay

5000 cells were seeded in each well of a 96-well plate containing 200 µl media and incubated overnight. The media was replaced with media containing varying concentrations of ADAADi derivatives. The cells were incubated at 37° C for the indicated time point. The media in each well was then replaced with media containing 0.45 mg/ml MTT and incubated for 2 hr at 37°C in a CO_2_ incubator. The media containing MTT solution was discarded and the purple precipitate formed was dissolved in 100 µl isopropanol. The plate was incubated for 15 min at room temperature and the absorbance was measured at 570 nm.

### Calculation of IC_50_ values

IC_50_ values were calculated using https://www.aatbio.com/tools/ic50-calculator/

### RNA extraction and cDNA preparation

RNA was extracted from untreated and treated HeLa cells as explained in Sharma *et al.* ^48^.

### qPCR

qPCR was done in 10 μl volume using 1X SYBR Green Master Mix,10 pmole of gene specific primers and 1μl of cDNA. The reaction was performed in an Applied Biosystems 7500 Real-Time PCR System using fast qPCR protocol (95°C, 20s; 1 cycle; 95°C, 3s; 60°C, 30s, 40 cycles).

### Western blotting

Cells were grown in 100 mm plates till 60-70% confluent. The cell extract was made by adding equal volume of RIPA buffer and freeze-thawing 3 times. The cell lysate was kept on rocker for 30 minutes at 4º C and clarified by centrifuging at 14000 rpm at 4º C for 10 min. The supernatant was transferred into a fresh tube and the protein concentration was determined using Bradford reagent. 100 µg protein was loaded into each well of a 10% SDS-PAGE. The protein bands were then transferred to PVDF membrane by electroblotting. After transfer, the blot was rinsed for 5 min each with 1X TBST five times and then incubated in 5% (w/v) skimmed milk in 1X TBST for 1 hour at room temperature. The membrane was again washed with 1X TBST five times for 5 minutes each and then incubated with primary antibodies with the required dilution overnight at 4º C. Subsequently, the membrane was washed five times for 5 minutes each with 1X TBST and then incubated for 45 min with secondary antibody conjugated with horseradish peroxidase. The membrane was washed 5 times for 5 min each with 1X TBST and finally rinsed once with 1X TBS. The blot was then developed by enhanced chemiluminescence method.

### PolyA mRNA purification

Poly(A) mRNA was purified using NEBNext® poly(A) mRNA magnetic isolation module (Catalog # <u>E7490S</u>), with certain optimization in the user protocol provided by the manufacturer. Briefly, 50 µl total RNA purified using Trizol (Sigma-Aldrich, USA) was added to 50 µl magnetic oligo d(T)25 beads and incubated at 65°C for 5 min and held at 4°C to denature the RNA. Subsequently, the beads were resuspended and incubated at room temperature for 5 min. This process was repeated twice and then the beads were washed with 200 μl of wash buffer provided by the manufacturer to remove the unbound RNA. The poly(A) mRNA beads complex was separated on magnetic stand, dried briefly at room temperature and resuspended in 17 μl of Tris buffer pH 7.5. The samples were incubated at 80°C for 2 min and then at 25°C to elute the poly(A) RNA from the beads. The purified poly(A) mRNA supernatant was transferred to nuclease free PCR tubes and the eluted poly(A) mRNA was quantified using NanoDrop™ 2000 spectrophotometer.

### RNA-seq library preparation

RNA-seq library was prepared using 100 ng purified poly(A) mRNA using NEBNext Ultra II RNA Library Prep Kit with certain optimization in the user protocol provided by the manufacturer. Briefly, the purified poly(A) mRNA was fragmented by mixing with NEBNext First Strand Synthesis Reaction Buffer (5X), Random Primers in 10 μl total volume and incubating the samples at 94°C for 7 min. After fragmentation, the samples first strand synthesis was performed using NEBNext First Strand Synthesis Enzyme Mix and incubating the samples at 25°C for 10 min, 42°C for 30 min and 70°C for 15 min. The second strand synthesis was performed on the same sample by using the NEBNext Second Strand Synthesis Enzyme Mix and buffer and incubating at 16°C for 60 min. After incubation, the samples were cleaned using 1.5X AMPure XP beads. The DNA bound to AMPure XP beads were separated using magnetic stand and washed three times with 80% ethanol. After drying, 20 μl of 0.1X TE pH 7.4 was added to the beads, mixed, and incubated at room temperature for 5 min. The eluted double strand DNA was proceeded for library prep using the same protocol as for the ChIP-seq library preparation.

### ChIP-seq library preparation

ChIP-seq library was prepared as per the instructions of user manual of NEBNext® Ultra™ DNA Library Prep Kit for Illumina® (Catalog # E7370) with certain optimization. Briefly, the library preparation was started with 100 ng of DNA purified after ChIP. The DNA was uniformly blunted for adaptor ligation using 3 μl End prep enzyme mix, 6.5 μl End prep 10X reaction buffer and nuclease-free water to final volume of 65 μl. The thermocycler was set with lid temperature 75°C and the samples were incubated at 20°C for 30 min and 65°C for 30 min. The samples were then transferred on to the ice and the components (15 μl Blunt ligase master mix, 2.5 μl NEB Next adaptor for Illumina, and 1 μl ligation enhancer to a total volume of 83.5 μl) for adaptor ligation were added directly into it. The reaction was mixed uniformly by pipetting and incubated at 20°C for 15 min. After incubation, 3 μl USER™ enzyme was added to the ligated samples, mixed, and incubated at 37°C for 15 min. After incubation, the samples were cleaned up with AMPure XP beads (0.9X). The DNA bound to AMPure XP beads were separated using magnetic stand and washed three times with 80% ethanol. After drying, 20 μl of 0.1X TE pH 7.4 was added to the beads, mixed, and incubated at room temperature for 5 min. The DNA was eluted (~15 μl) and mixed with 25 μl NEBNext Q5 Hot Start HiFi PCR Master Mix, 5 μl Index Primer/i7 Primer, and 5 μl Universal PCR Primer/i5 Primer. The samples were uniformly mixed, and PCR was performed (initial denaturation at 98°C for 30 sec, 10 cycles of cycling phase of 98°C for 10 sec and 65oC for 75 sec and final extension at 65°C for 5 min). After PCR, the samples were cleaned up using AMPure XP beads (0.9X). The DNA bound to AMPure XP beads were separated using magnetic stand and washed three times with 80% ethanol. After air drying, 20 μl of 0.1X TE was added to the beads, mixed and incubated at room temperature. The DNA (~15 μl) was eluted and the concentration was checked using NanoDrop™ 2000 spectrophotometer. The sequencing was done by Genotypic Technologies, Bengaluru, India

### RNA-seq and ChIP-seq data analysis

Quality of the RNA-seq data was assessed by FASTQC analysis. Reads were quasi-aligned onto the reference human transcriptome using salmon to obtain the transcript counts. Salmon generated transcript quantification were converted to gene level quantification using tximport (version 1.10.1) package and imported to DESeq2 (version 1.22.2) on R (version 3.5.2) for gene level analysis. Normalisation and differential gene expression analysis was done using DESeq2 and Differential transcript usage analysis was done by rnaseqDTU package. For Chip-Seq, raw reads were trimmed by trimmomatic (version-0.36.5) to remove adaptor sequence and base quality was checked by FASTQC. Trimmed reads were aligned to human genome (hg38) by BOWTIE2 with default settings. Aligned files were marked for duplicates by PicardMarkduplicates and filtered on bit wise flags by SAM tools on Galaxy platform. Only paired end reads that are mapped in proper pair were selected for peak calling. Biological replicates of SMARCAL1 and BRG1 were merged in a single BAM file before peak calling. Peak calling was performed by MACS2 (Version 2.1.1.20160309.0) with default settings. Both alignment and peak calling was done on the galaxy platform (https://usegalaxy.org). Gene annotation was done using chipseeker package on R.

### Comet assay

Single cell gel electrophoresis was performed as described by Nandhakumar *et al*. ^49^. Briefly, HeLa cells were grown till they were 60-70% confluent and then given treatment with 5 µM ADAADiN for the desired time period. After appropriate treatment, the cells were washed in cold 1X PBS, trypsinized and resuspended in ice-cold PBS. The cell suspension containing 10^4^ cells/slide was embedded in 100□µl of 1% low-melting agarose (Sigma-Aldrich, USA) in 1X PBS and spread onto microscopy slides coated with 1% normal-melting agarose. The cells were lysed in the lysis solution (2.5□M NaCl, 100□mM EDTA, 10□mM Tris base, 8□g/L NaOH to adjust pH 10; 1% Triton X-100, and 10% DMSO) at 4°C for 2 hr. Following lysis, the slides were placed for 20□min in a tank with cold electrophoresis buffer (300□mM NaOH, 1□mM Na_2_EDTA, pH 12.5) and electrophoresed for 30□min at 25□V and 300 mA. The slides were then neutralized with 0.4□M Tris-Cl pH 7.5 and the DNA was stained with 0.5 mg/ml ethidium bromide. The slides were analyzed using fluorescence microscope (Nikon) at 10X magnification. Casplab software was used for the analysis of tail moment according to the given formula which is a measure of both the smallest detectable size of migrating DNA (represented by comet tail length) and the number of relaxed / broken pieces (represented by DNA intensity in the tail) ^50^.

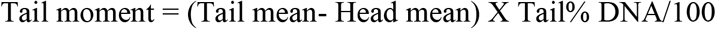

### FACS analysis

Cell were grown in 35 mm dishes till they were 60-70% confluent and then given treatment with ADAADiN for the desired time period. After rinsing with 1 ml of chilled 1X PBS, the cells were trypsinized and collected in 1 ml of chilled 1X PBS. The cell pellet obtained by centrifuging at 1200 rpm at 4º C for 10 min was resuspended in 70% ethanol gently using vortex to avoid clumping and incubated for 30 min at 4ºC. The cells were washed twice with 1 ml of chilled 1X PBS at 1200 rpm at 4º C for 10 min. Subsequently, the cells were treated with 50 μl of 100 μg/ml stock of RNaseI and then incubated with 200 μl of 50 μg/ml propidium iodide for 45 min at room temperature prior to analysis. Analysis was done using BD FACS Calibur 4C and Modfit software.

### Immunofluorescence

Cells (HeLa and DU145) were seeded and grown till they were 60-70% confluent and then given treatment with sub-lethal concentration of ADAADiN for desired time period. The cells were fixed in methanol: acetone (1:1), permeablized with 0.5% Triton X-100, washed and blocked with 1% BSA in 1X PBS overnight at 4°C. The cells were washed with 1X PBS and d and incubated with γH2AX antibody (1:120 dilution) for 60 min at room temperature. The cells were washed with 1XPBS and then incubated with secondary antibody solution (1:200 dilution) and Hoechst (1:100 dilution) at room temperature for 60 min. The cells were washed with1 ml of 1XPBS five times, covered with a coverslip using 70% glycerol and analyzed using confocal microscope (Nikon TiE) under a 60X oil immersion objective.

### *In vivo* ATPase activity analysis

HeLa cells were grown in 100 mm dishes till they were 60-70% confluent and then treated with 5 µM ADAADiN for the desired time period. The cells were trypsinized and washed with 1X PBS twice followed by incubation on rocker at 4°C for 1 hour in 200 µl lysis buffer (50 mM Tris-Cl (pH 7.5), 400 mM NaCl, 1 mM EDTA, 1 mM EGTA, 0.1% NP-40, 1 mM PMSF, and protease inhibitor cocktail) and then sonicated in water bath (4 cycles; 10 s ON and 50 s OFF). The cell lysate was clarified by centrifuging at 13,000 rpm for 10 min at 4°C. The lysate was pre-cleared using 20 µl protein G beads and the pre-cleared supernatant was incubated overnight at 4°C with 1 µl polyclonal antibodies either against SMARCAL1 or BRG1. 50 µl equilibrated Protein G beads were added to immunoprecipitated protein bound to the polyclonal antibodies. The beads were centrifuged at 2500 rpm for 3 min, the supernatant was discarded, and the beads were washed with lysis buffer four times. The ATPase activity of the immunoprecipitated SMARCAL1 and BRG1 was estimated using NADH oxidation assay at 340 nm as detailed in Dutta *et al*. ^6^.

### Acridine Orange/Ethidium bromide staining to quantitate apoptotic cells

The number of apoptotic cells after ADAADiN treatment were quantitated using Acridine Orange/Ethidium bromide staining as per the protocol described by Ribble *et al.* ^51^. Quantitation and analysis was done as described by Anasamy *et al*. ^37^.

### Colony formation assay

DMEM (2X) containing FBS and antibiotics was mixed with equal volume of agar (1% w/v) at 42°C. To prepare an agar bed, a 6-well plate was plated with 1.5 ml of the DMEM-agar mix and allowed to solidify. HeLa and DU145 cells (either untreated or treated with sub-lethal concentration of ADAADiN for 24 hr) were trypisinised and the pellet was mixed with 5 ml of 2X DMEM and 5 ml of 0.6% (w/v) agar. This cell mix (1.5 ml) was plated on top of the agar bed. After a short incubation at room temperature that allowed the gel to solidify and form a mesh around the cells, 1.5 ml of 1X DMEM containing FBS and antibiotics was added. To study the effect of ADAADiN, sub-lethal concentration of ADAADiN was added to the media. The plates were incubated in the CO_2_ incubator at 37°C and images were taken on the 4^th^, 8^th^, 12^th^ and 16^th^ day using Nikon TiS microscope at 5X, 20X and 40X magnification.

### Wound healing assay

HeLa and DU145 cells were grown in 35 mm dish to 80% confluency before adding sub-lethal concentration of ADAADiN for 24 and 48 hr. A scratch was made with 10 μl pipette tip in the middle of the dish. The media was aspirated off to remove the dead cells and fresh media was added. The media for the control cells did not contain ADAADiN while in the ADAADiN treated cells, the fresh media contained sub-lethal concentration of ADAADiN. The migration of the cells and the recovery of the wound at every 12 hr time point were monitored using Nikon microscope TiS at 5X magnification.

### Annexin V-FITC apoptosis detection

The cells were grown in 60 mm dish to 60-70% confluency before treatment with sub-lethal concentration of ADAADiN for desired period of time. The cells stained with Annexin V and PI using Annexin V-FITC apoptosis detection kit as per the manufacturer’s instructions (eBiosciences, USA) (Catalog # BMS500FI/100). The samples were analyzed in BD FACS Calibur 4C flow cytometer.

### Zymography assay

To assess the activity of metalloproteinases, zymography assay was performed. Cells (untreated and ADAADiN treated) were grown in serum-free DMEM media for 12 hr and 50 μl of the media was loaded onto an 8% SDS-polyacrylamide gel containing 0.2% (w/v) gelatin.

The samples were electrophoresed at constant voltage (125 V) till the dye front reached the end of the gel. The gel was renatured by incubating it in buffer containing 50 mM Tris-Cl, pH 7.5, 5 mM CaCl_2_, 1 μM ZnCl_2_, 2% (w/v) NaN_3_ and 2.5% (v/v) Triton-X100 for 60 min at room temperature. The renatured gel was then transferred to incubation buffer (50 mM Tris-Cl, pH 7.5, 5 mM CaCl_2_, 1 μM ZnCl_2_, 2% (w/v) NaN_3_ and 1% (v/v) Triton-X100) for 24 hr. After incubation, gel was washed and stained with Coomassie Brilliant Blue R-250 for 1 hr and distained in destaining solution. The breakdown of gelatin was monitored as the appearance of white band in the background of blue stain.

## AUTHOR CONTRIBUTIONS

Conceptualization, R.M., J.W.H., R.R., S.H., U.B.C.; Methodology, R.M. R.R., S.H., K.G., D.B., and U.B.C.; Investigation, R.R., S.H., K.G., D.B., S.S., U.B.C.,; Resources-J.W.H. and R.M.; Writing-original draft, R.M. and R.R.; Writing-review and editing, R.M., J.W.H., R.R., and S.H. Funding acquisition, R.M.; Supervision, R.M.

## ACKNOWLEDGEMENTS

The authors would like to thank the Central Instrumentation facility, School of Life Sciences for confocal microscope and FACS facility. The authors would also like to thank Dr. Sarika Gupta for technical support.

