## Supplementary File for "Altering Mammalian Transcription Networking with Adaadi: An Inhibitor of Atp-Dependent Chromatin Remodeling"

**Supplementary Figure Legends**

**Supplementary Figure 1. ADAADi induces cell death in cancer cell lines**. Kill curves showing the effect of ADAADi after treatment. (A). HeLa (B). DU145 (C). MCF-7 (D). L132 (E). PC3 (F). MDA-MB-231 (G). A549 (H) HepG2 (I) HEK293.

**Supplementary Figure 2. (A).** The counts per sample before and after normalization. (B) Mean-variance relationship between the untreated and ADAADiN treated RNA-seq samples. (C). Correlation between untreated and ADAADiN treated RNA-seq samples. (D). Representative chip-seq peaks visualized using IGV genome browser showing occupancy of SMARCAL1 and BRG1on *ABCG1, ABCB1* and *ABCC3* genes.

**Supplementary Figure 3**. Comparison of BRG1 ChIP-seq data with the publicly available dataset. Representative chip-seq peaks visualized using Easeq showing occupancy of BRG1 on *SMARCAL1, SMARCA4 (BRG1), DORSHA, DICER, ATM, ROCK2, EGF, MALAT1, MMP2,* and *MMP9* genes. In each of the figure, A represents the ChIP performed with BRG1 in this paper, B represents the Input sample for this experiment, C represents the ChIP dataset available in the public domain and D represents the Input sample for the dataset available in the public domain. The green box indicates the presence of BRG1 on the gene that is common to both the datasets.

**Supplementary Table 1: List of primers used for qPCR analysis.**

| **Gene** | **Forward primer (5'→3')** | **Reverse primer (5'→3')** |
| --- | --- | --- |
| *GAPDH* | GGTCGGAGTCAACGGATTTGGTC | GAGGGATCTCGCTCCTGGAAG |
| *BRG1* | GCGAGTACAGGCTGCAGGCT | TGGCGCAGCTGCCTCTG |
| *SMARCAL1*  (used for HeLa) | TCCCATCTGTTCATTGAATATATCTTGGAC | GCTGCACGTGCTTTCTCTTCAAGCTC |
| *SMARCAL1*  (used for DU145) | CACCAAGGACAAAACTAAACAG | GTCCAAGATATATTCAATGACAGATG |
| *ATM* | GGGCAGTCACGCAGGGTTTG | ATCACTGTCACTGCACTCGGAAGG |
| *BRCA1* | ACCCTCTGCTCTGGGTAAAGTTCA | TGGTCACACTTTGTGGAGACAGGT |
| *MMP-2* | GATGTCGCCCCCAAAACGGAC | CACCTGTCTGGGGCAGTCCAAA |
| *MMP-9* | GTGCCGGAGGCGCTCATGTA | GGCTCAGGTTCAGGGCGAGG |
| *CDH1* | GCTGTGTCATCCAACGGGAA | CACCTTCCATGACAGACCCC |
| *VIM* | GGCTCGTCACCTTCGTGAAT | CAGAGAAATCCTGCTCTCCTCG |
| *CDH2* | GGGTCATCCCTCCAATCAAC | ACCTGATCCTGACAAGCTCT |
| *FN1* | GGCTGTCAGTCAAAGCAAGC | TCGCAGTTAAAACCTCGGCT |
| *CEP135* | TCCTTCAAGTGGCTGATAAC | GCAACTGACAGTCGTTCTAT |
| *CENPE* | CTTGGGAGGAAATGCAAAGA | GGAGAGCTTCATCAGTTGATAC |
| *KMT2A* | GAGAGGATGAGCAATTCTTAGG | ACAGATGGATCTGAGAGGATAG |
| *NEK1* | GGACAAGAAGGAAGTGAAGAG | ACTCCTTTAGCCACCAAATG |
| *ROCK2* | CTTGCTGGATGGCTTAAATTC | ATCATAGTCTTCTGCCTTCATC |
| *DCTN3* | GGACAGTGCTCACATCAAA | ACGAATTGCTTGGAGAGAAG |
| *SNAPC2* | GCTGTGGACTTTGAGAAGAT | GTAGGTACGTCTCCGTCATA |
| *EMP3* | TGTCTCTCCTTCATCCTGTTC | TAGATCAAGGCGCCAGTAAA |
| *MALAT1* | CATTCCAGGTGGTGGTATTT | TTCTGTGTTATGCCTGGTTAG |
| *EGF* | GGAGGTTCTGTCCACATTAG | CCTTTCCAGTGTGTTTGTTG |
