## Supplementary figures and images for "Altering Mammalian Transcription Networking with Adaadi: An Inhibitor of Atp-Dependent Chromatin Remodeling"

### Supplementary Figure 1

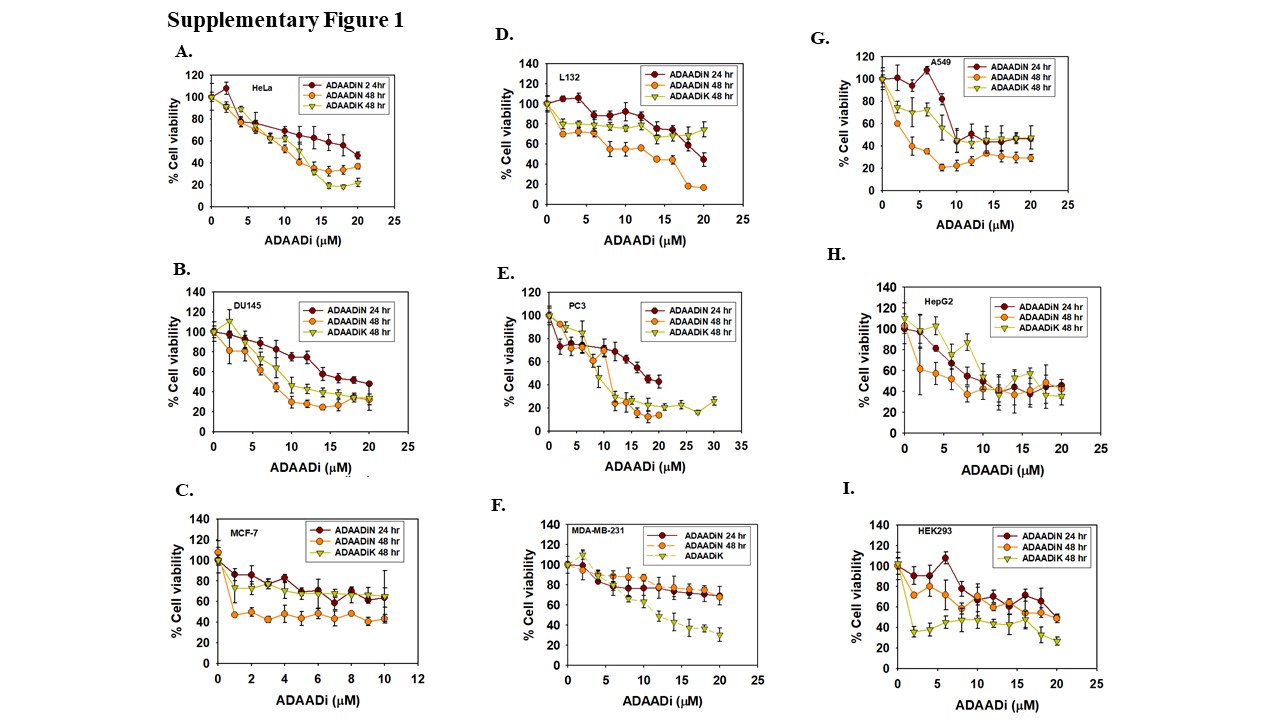

### Supplementary Figure 2

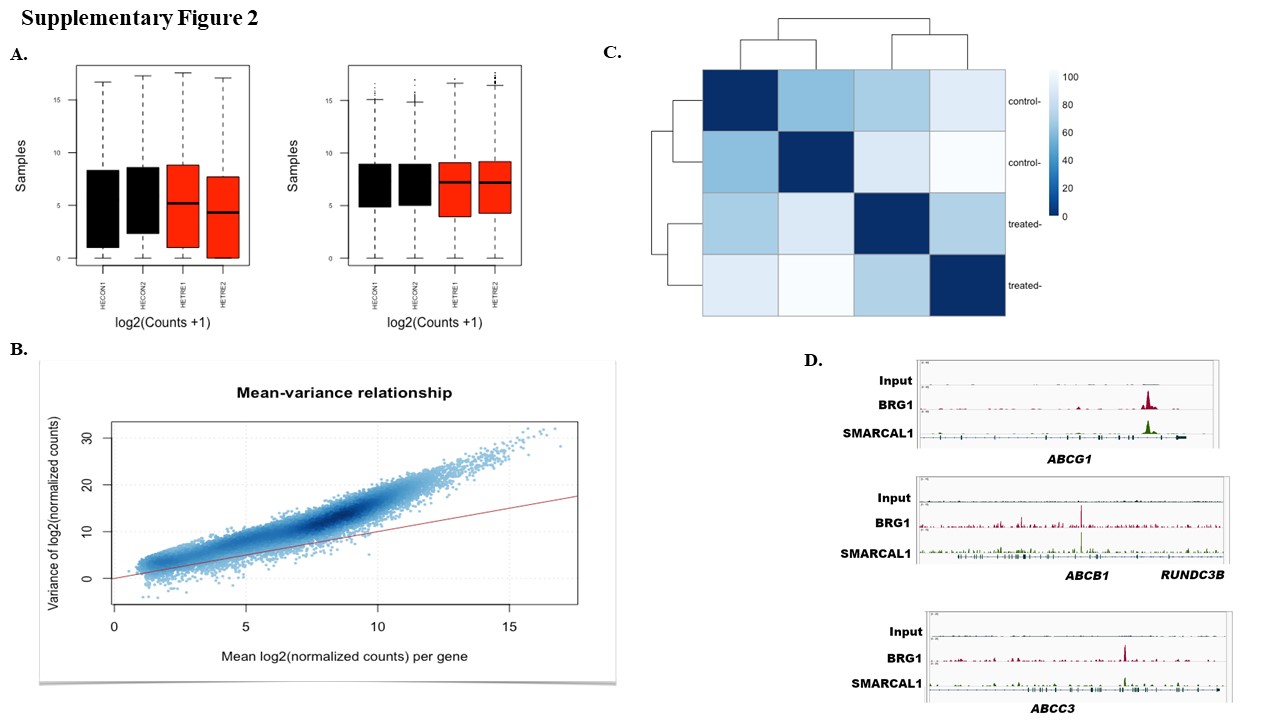

### Supplementary Figure 3

#### Supplementary Figure 3

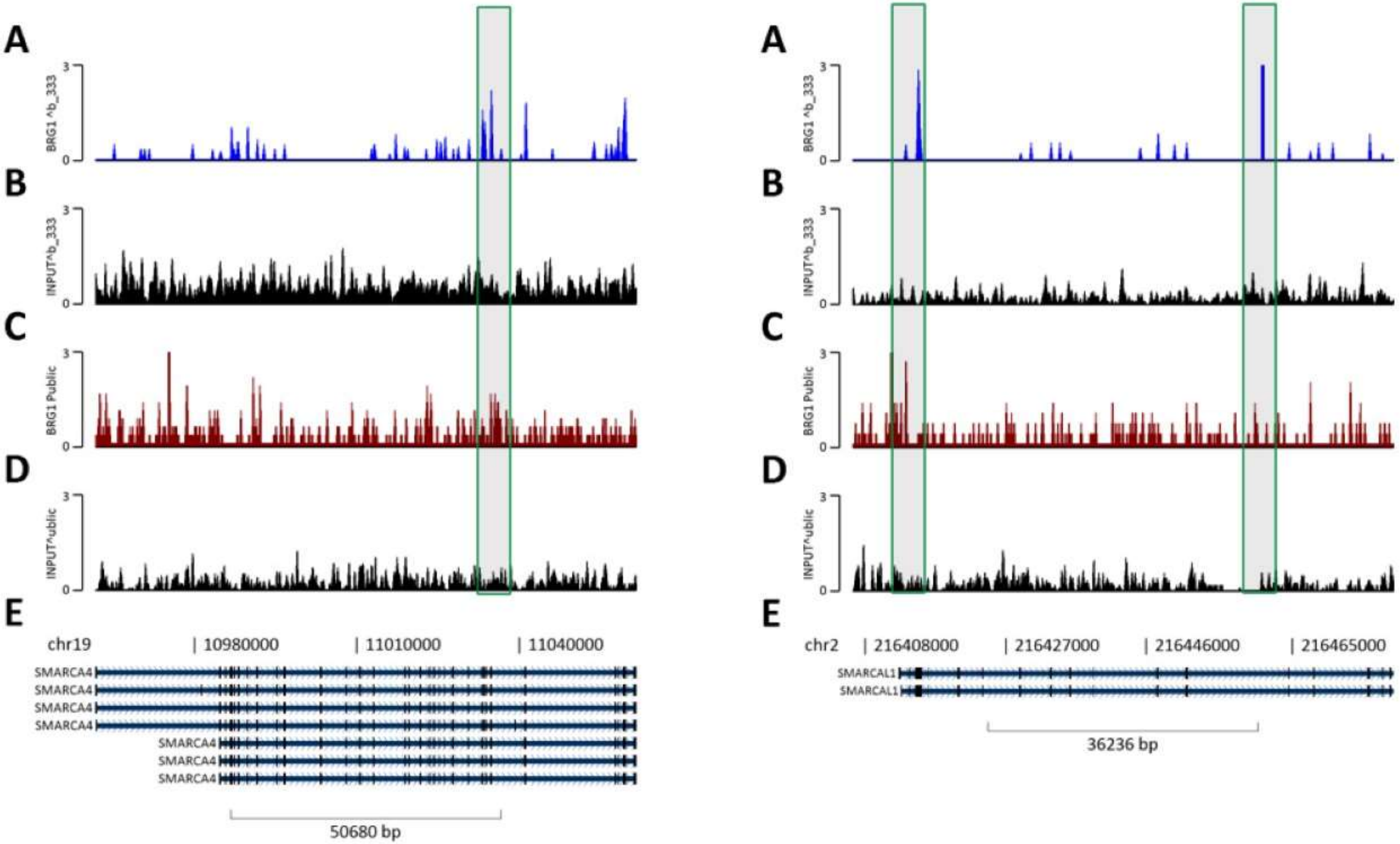

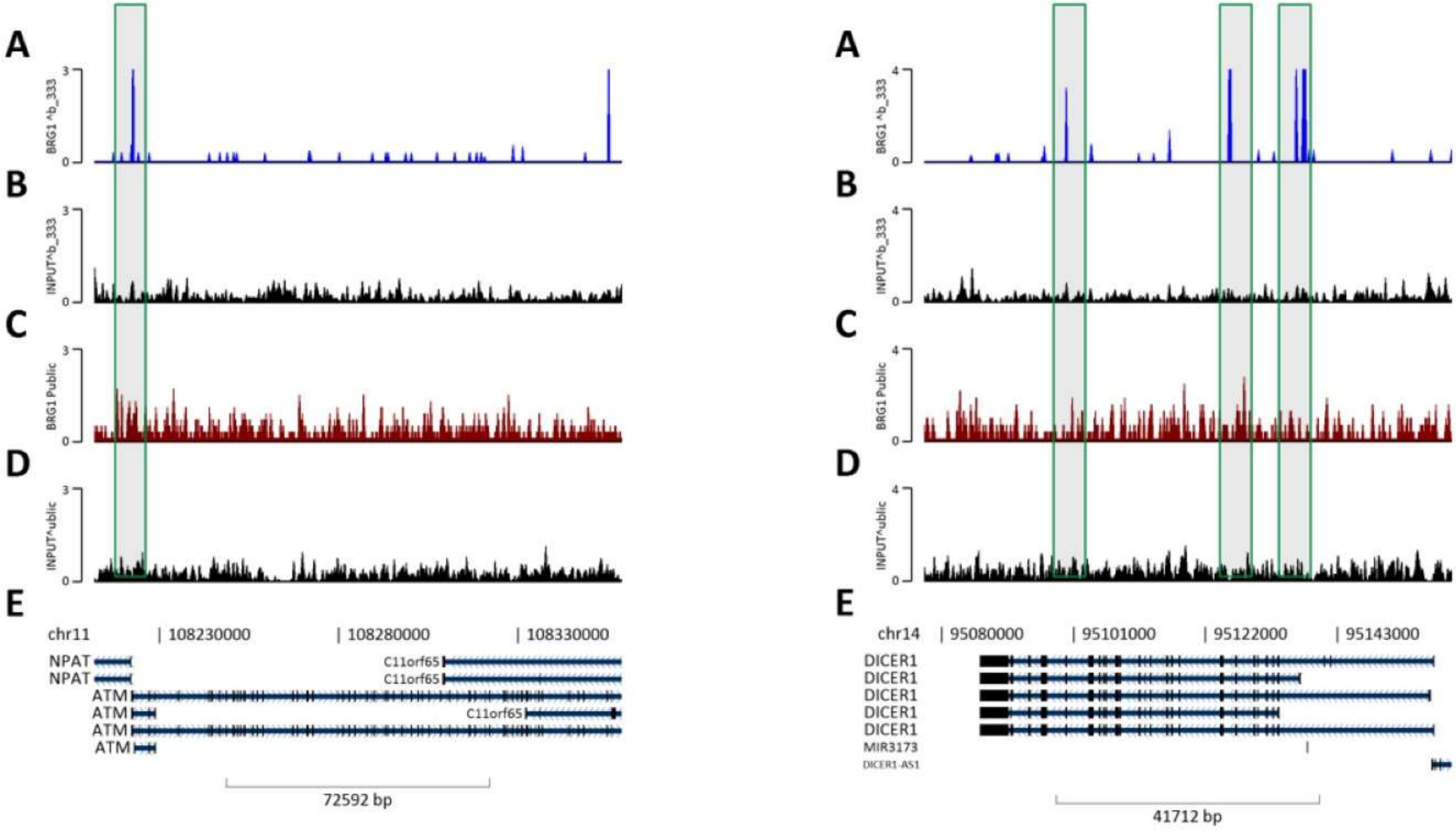

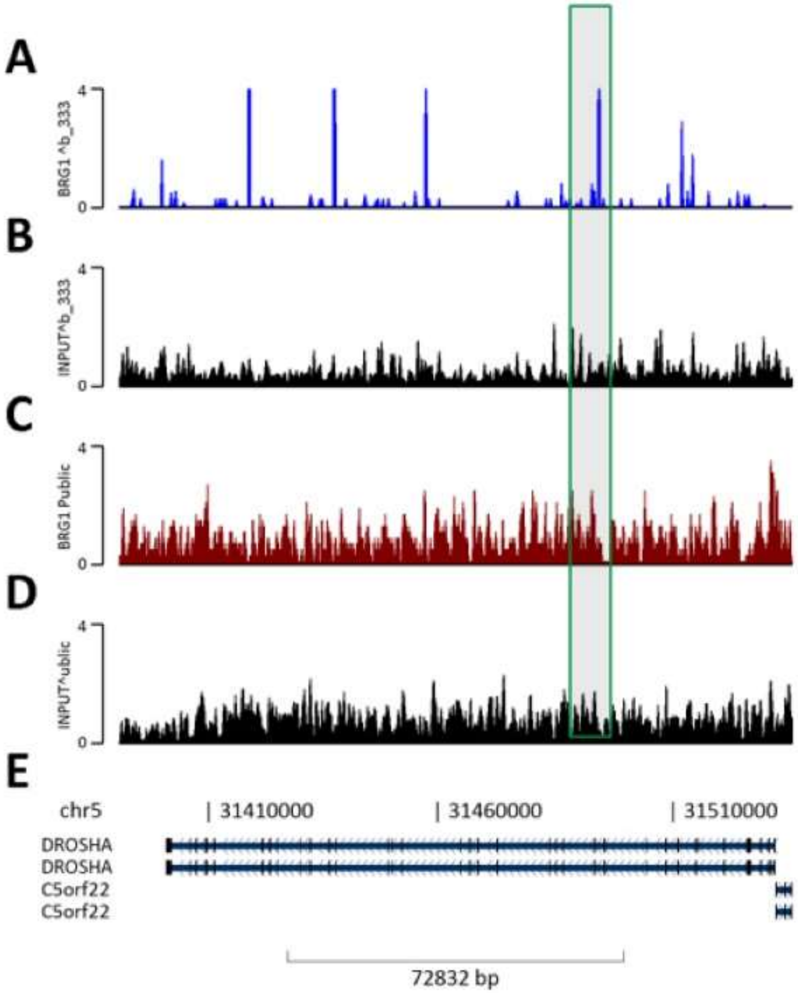

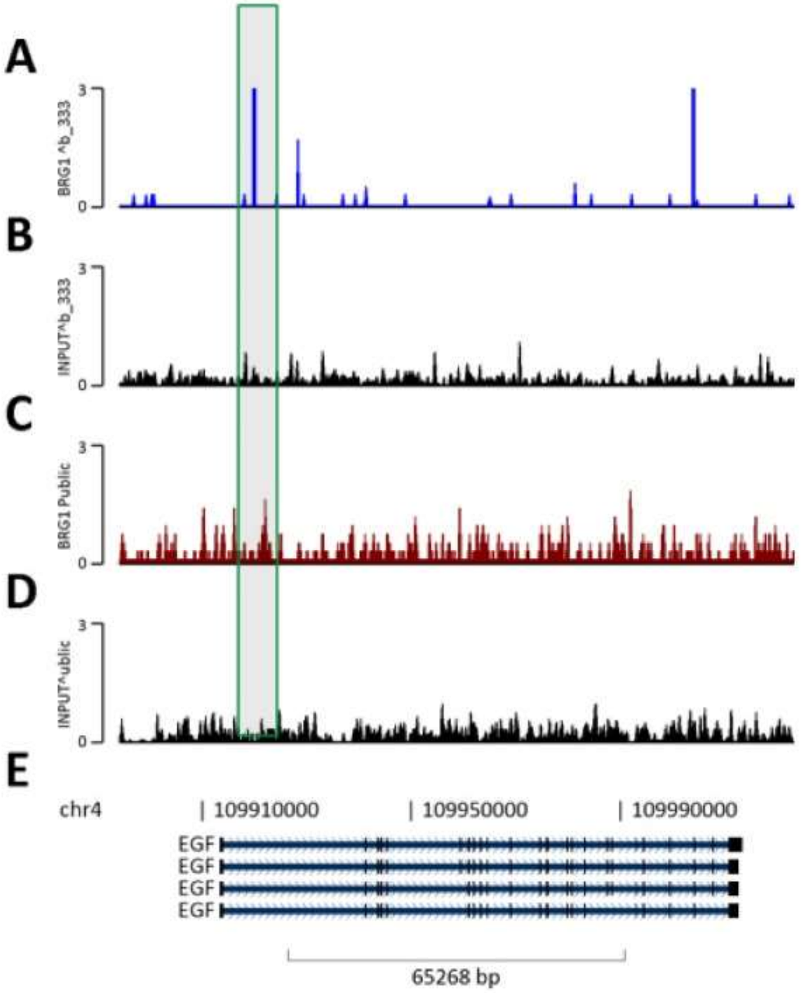

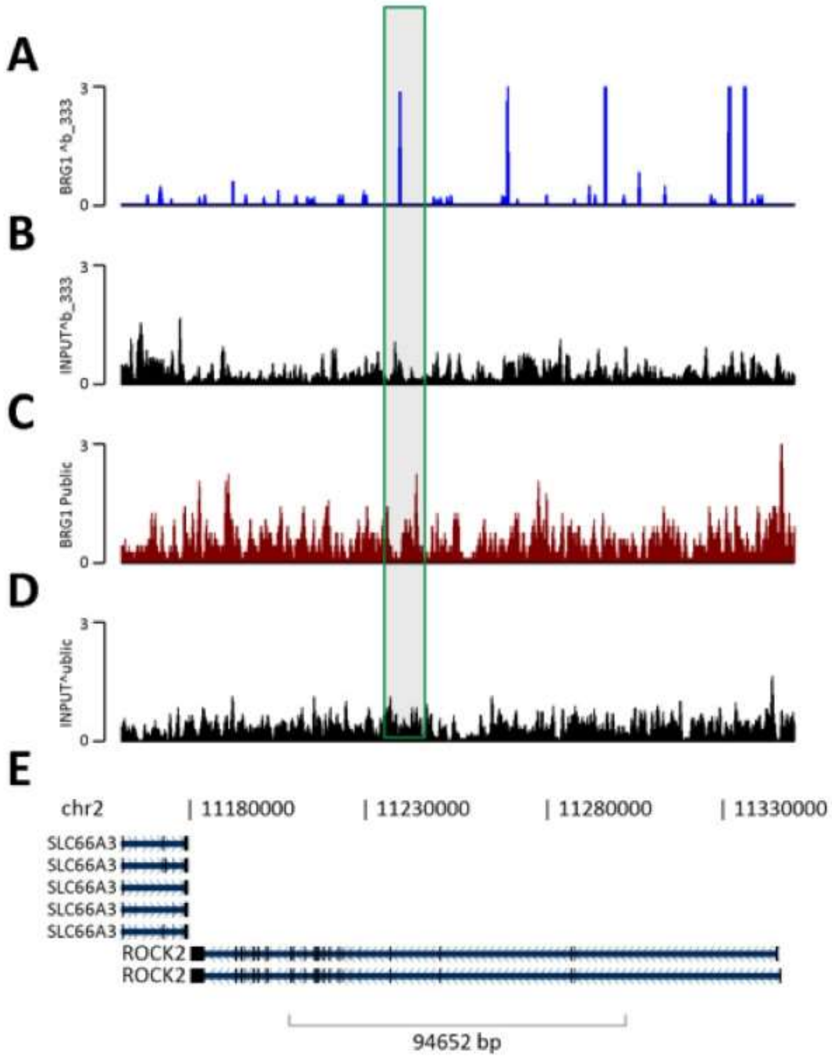

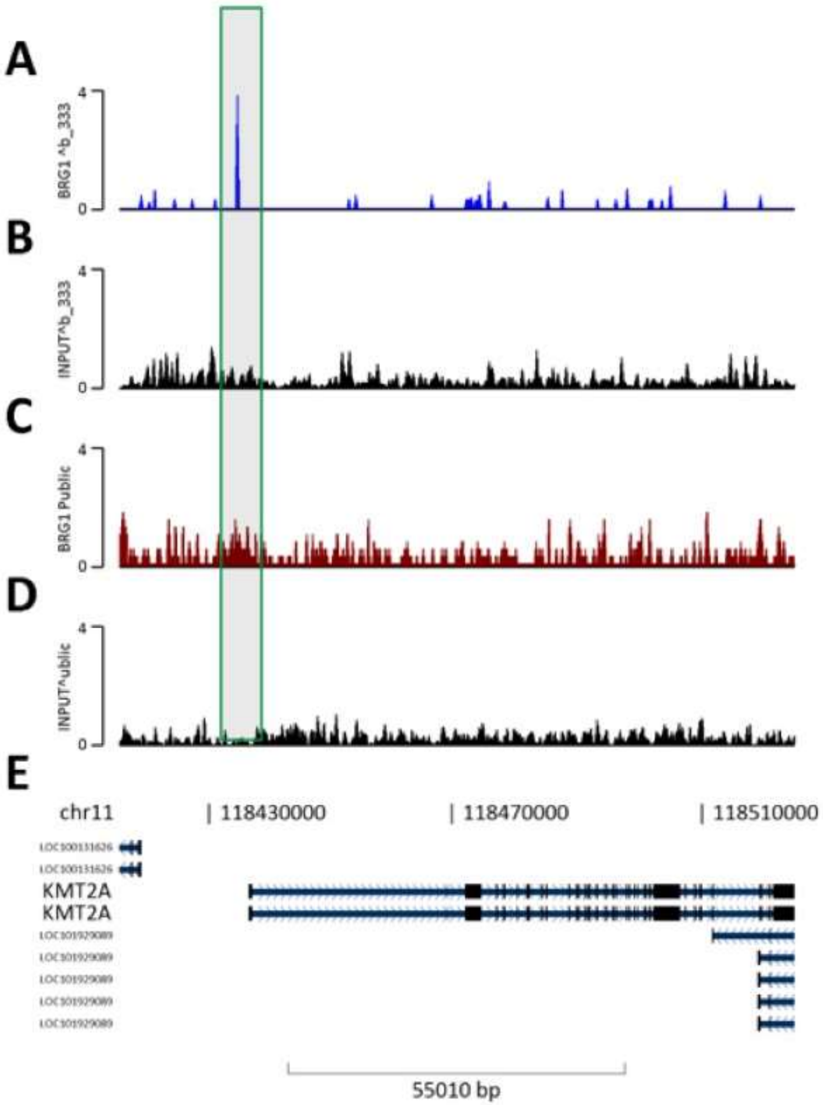


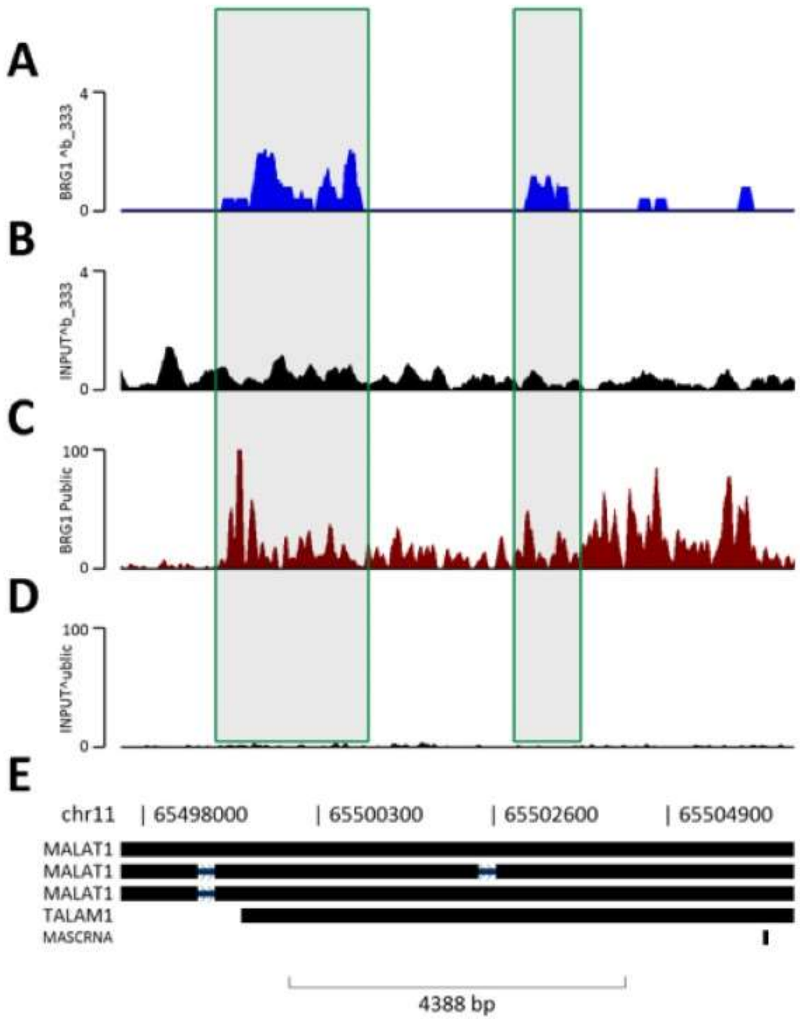


**Here Scale diff.

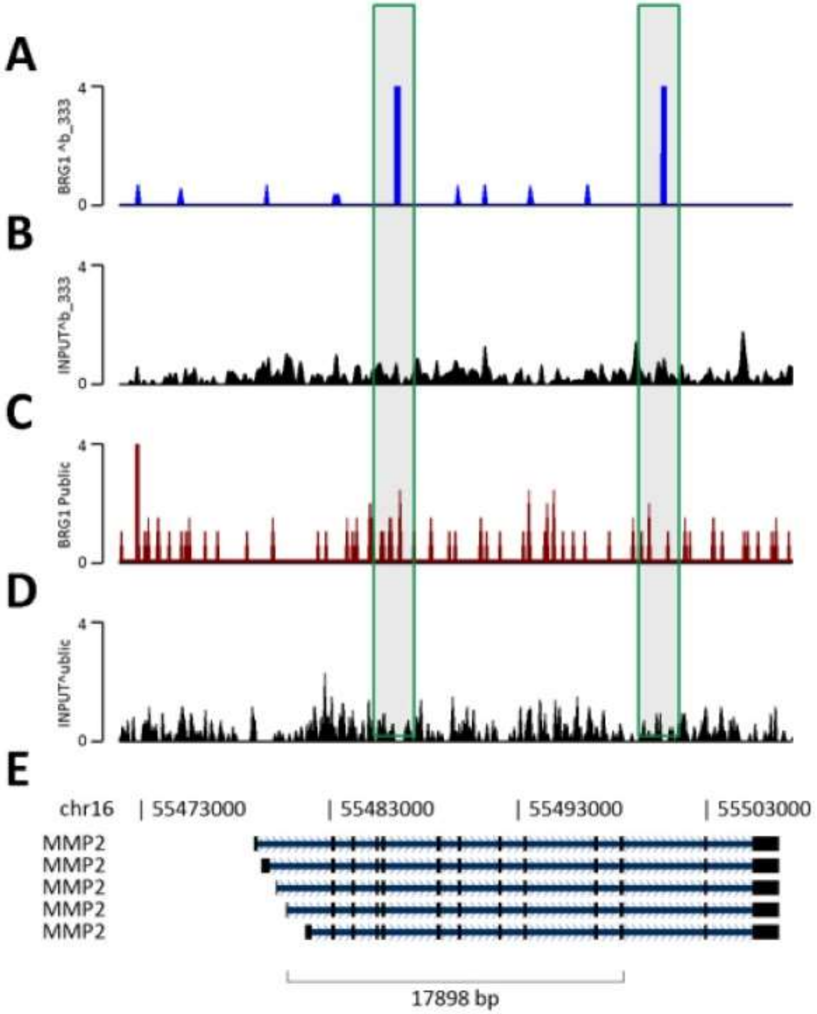

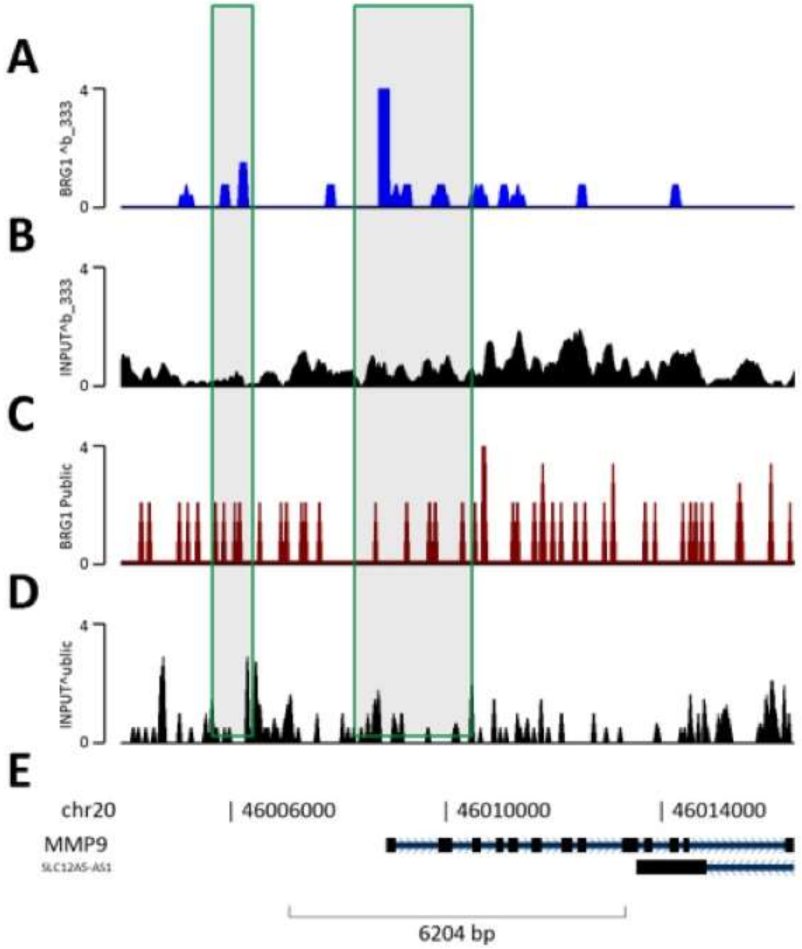

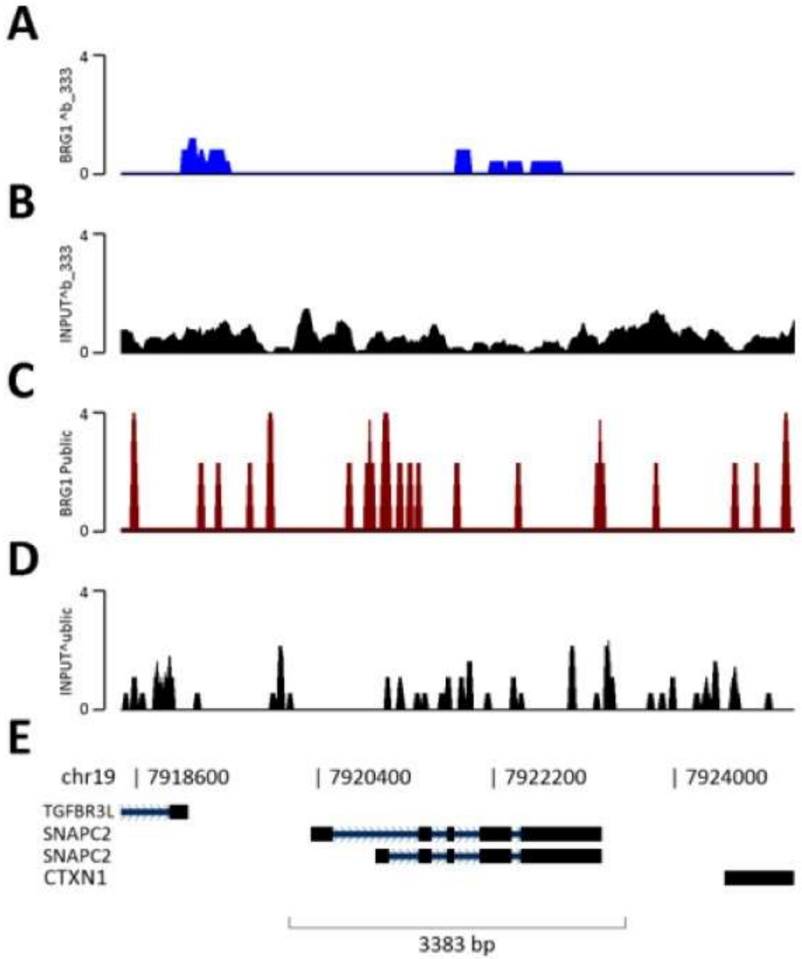
